# Computational optical sectioning by phase-space imaging with an incoherent multiscale scattering model

**DOI:** 10.1101/2020.11.29.402958

**Authors:** Yi Zhang, Zhi Lu, Jiamin Wu, Xing Lin, Dong Jiang, Yeyi Cai, Jiachen Xie, Tianyi Zhu, Xiangyang Ji, Qionghai Dai

## Abstract

Optical sectioning is essential for fluorescence imaging in thick tissue to extract in-focus information from noisy background. Traditional methods achieve optical sectioning by rejecting the out-of-focus photons at a cost of photon efficiency, resulting in a tradeoff between sectioning capability and detection parallelization. Here, we show phase-space imaging with an incoherent multiscale scattering model can achieve computational optical sectioning with ~20 dB improvement for signal-to-background ratio in scattering medium, while maximizing the detection parallelization by imaging the entire volume simultaneously. We validated the superior performance by imaging various biological dynamics in *Drosophila* embryos, zebrafish larvae, and mice.

---

The beauty of life lies in the complexity and variety of cellular behaviors in three-dimensional (3D) living organisms, which are difficult to appreciate without optical sectioning^1–3^, as the information can be easily flooded in the severe noisy background. Various methods have been proposed to achieve optical sectioning by rejecting the out-of-focus photons either physically or computationally, such as confocal microscopy^4,5^, two-photon microscopy^6,7^, light-sheet microscopy^8,9^, and structured illumination microscopy^10,11^. However, these methods still require scanning of points, lines, or planes for 3D imaging, leading to the tradeoff between the sectioning capability and detection parallelization^1,3,12^. Such a tradeoff intrinsically restricts the 3D imaging speed in fluorescence imaging, especially with a limited photon budget set by the sample health^1,12,13^. In addition, it’s hard to remove the scattered photons due to the lack of depth-dependent features.

Phase-space imaging, by collecting the high-dimensional local variances of the coherence, provides a new imaging framework with digital synthesis of the partially-coherent light field, which shows strong robustness to scattering and aberrations^14–18^. As a typical example, light-field microscopy (LFM)^19^ captures the phase-space measurements within a snapshot by a microlens array^20^, facilitating various applications in biology, such as imaging hemodynamics^21^ and large-scale neural activities^14,22^. Different from wide-field imaging with a shallow depth of field (DOF)^23–25^, LFM keeps the photons focused along different angles within the extended depth of field and maximize the parallelization by imaging the entire volume simultaneously. However, it suffers from great degradation in thick tissue such as the mammalian brain, due to scattering and dense fluorescence labeling. While many efforts in hardware have been made to achieve additional optical sectioning^26–28^ at the cost of temporal resolution, space constraints and system compactness, the imaging model of this computational framework has barely been explored, especially for complicated imaging environment in deep tissue^29,30^.

Here, we show the necessity of an accurate imaging model in the complete space to unlock the full power of phase-space imaging in thick tissue. With an incoherent multiscale multiple-scattering model, phase-space imaging can achieve snapshot quantitative 3D information with optical sectioning computationally in densely labeled or scattering samples. This method is termed as quantitative LFM (QLFM). We find that the severe degradation of traditional LFM in thick tissue mainly results from the incomplete space and ideal imaging model utilized during reconstruction^14,29,30^. By building up the incoherent imaging process in phase-space domain accurately with various factors, including the nonuniform resolution of different axial planes, out-of-focus fluorescence across a large range, multiple scattering, and system aberrations, we can not only improve the resolution and contrast with significantly-reduced computational costs, but also achieve two-orders-of-magnitude improvement in signal-to-background ratio (SBR) over traditional algorithms and wide-field microscopy (WFM), which is critical for quantitative biological analysis. To demonstrate the versatile applications, we imaged various 3D biological dynamics in different specimens, including the zebrafish larvae heart beating, blood flow, and whole-brain neural activity, *Drosophila* embryo development, and mouse brain neural activity with notably better performance than that of traditional ideal models without increasing the system complexity.

## Results

### Principle and implementation of QLFM

As a general computational model for incoherent conditions, our QLFM method is compatible with different schematics of phase-space imaging. Here, we chose the unfocused LFM for experimental demonstrations by inserting a microlens array at the image plane of a normal wide-field microscope (Fig. 1a). A 4f system was used to relay the back focal plane of the microlens array at the image sensor, with each microlens covering approximately 13 × 13 sensor pixels (Methods). Once set up, the high-resolution 3D range of LFM was fixed with a specific objective. To flexibly adjust the volume size for different specimens, we applied a piezo stage for high-speed axial scanning at large axial steps^9^. With the matched numerical aperture (NA), LFM provides an extended depth of field ~169-times larger than that of WFM, which makes it very different from other wide-field imaging methods. The fluorescence signals far from the focal plane exhibit similar Gaussian backgrounds in WFM^23–25^, while they show apparent distinguishable features between different angular components captured by LFM within a snapshot (Supplementary Fig. 1 and Supplementary Video 1). We find that these depth-dependent features provide LFM with a similar capability of computational optical sectioning as structured illumination microscopy^31^. The 3D out-of-focus fluorescence up to the centimeter scale can be reconstructed at low resolution, as long as we take it into consideration in the multislice model as a complete space. These layers far from the native objective plane have usually been neglected in previous methods due to the limited computational resources and contributed considerable background noises and artifacts in the high-resolution range (Fig. 1b). To address this problem, we propose a multiscale model in the phase-space domain by resampling the volume based on the nonuniform resolution of different axial planes, avoiding many unnecessary calculations in a complete space (Fig. 1b, Supplementary Fig. 2 and Methods). We then conducted a numerical simulation with gradually increasing background levels to show the significantly improved quantitative property of the multiscale model, which is barely considered in previous studies but important for biological analysis (Supplementary Fig. 3).

**Fig. 1.**
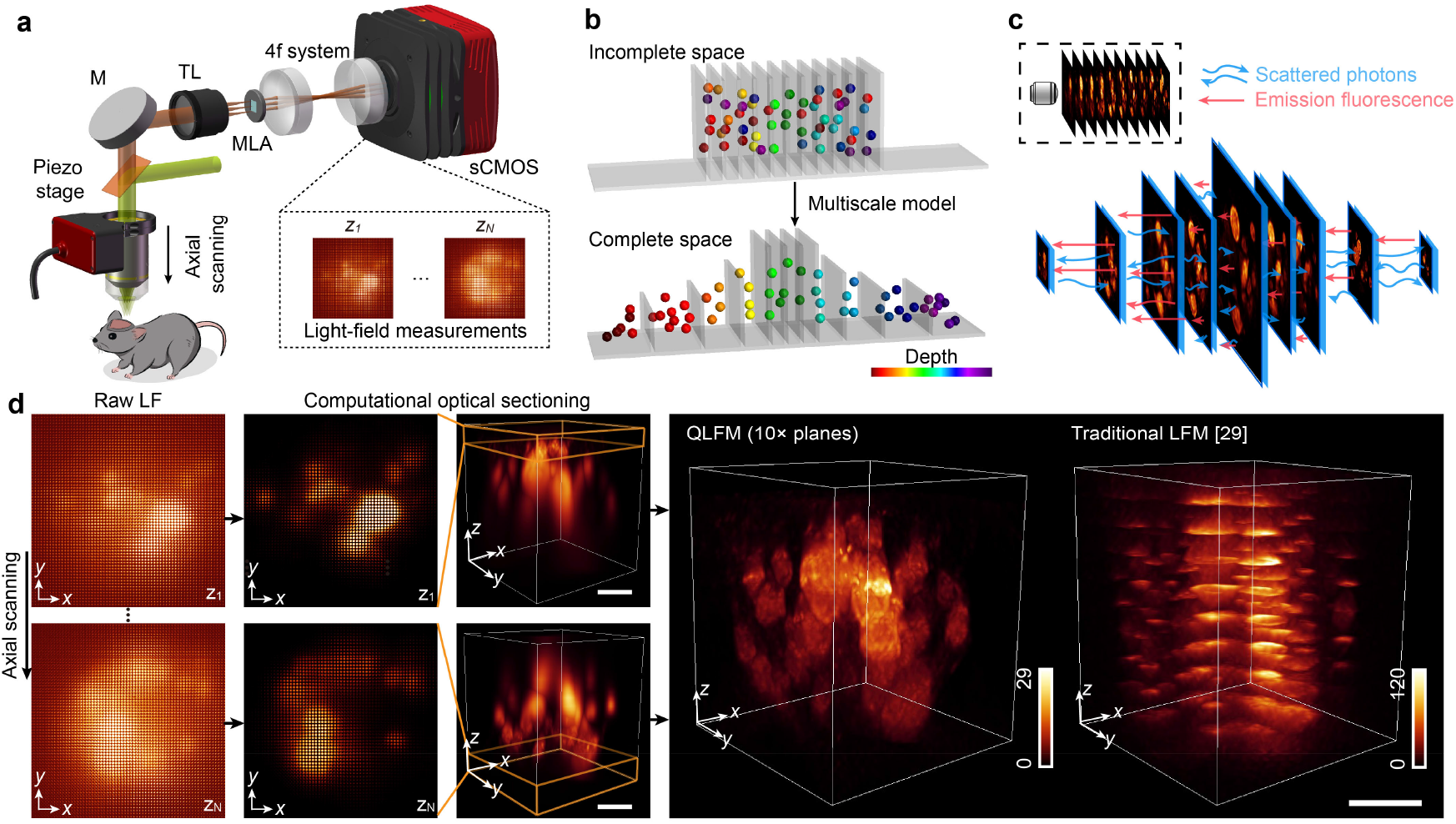
Schematic of quantitative light-field microscopy (QLFM). **a**, Experimental setup of our QLFM system with a simple microlens array (MLA) inserted at the image plane for snapshot phase-space measurements. A piezo stage is used for high-speed axial scanning at a large step size periodically. **b**, Concept of the multiscale model. Incomplete space used in traditional LFM reconstruction results in strong background noise and loss of quantitative properties in complicated environments. We apply different sampling rates in 3D based on the effective resolution of LFM at different axial planes to model a large volume for background rejection with an orders-of-magnitude reduction in computational cost. **c**, Schematic of the multiscale scattering model. We differentiate the multiple-scattered photons from native emission fluorescence in the multislice model based on the first Born approximation to retrieve 3D fluorescence quantitatively in deep tissue. **d**, Illustrations of the algorithm by imaging GFP-labeled B16 tumor spheroids. Due to the extended depth of field, even the out-of-focus fluorescence far from the native objective plane has apparent depth-dependent features in LFM, which can be reconstructed in the multiscale model for computational optical sectioning. The axially scanned LF images of 10 planes can be realigned into multiple angular focal stacks to reconstruct the entire volume as a whole without artifacts. QLFM shows greatly improved resolution and contrast over traditional LFM which reconstructs each subvolume separately. Scale bars, 30 μm.

Except for out-of-focus fluorescence, the scattered photons pose another challenge in thick tissue, which cannot be rejected by optical sectioning due to the depth-independent property. Although multislice scattering models have achieved great success in the coherent imaging modalities^32^, they are barely considered in deconvolution algorithms for incoherent fluorescence imaging. As the 4D phase-space measurements can fully describe the partially coherent light-field distributions, they provide an opportunity to infer the native 3D fluorescence as well as the nonuniform 3D scattering coefficients. Here, we derived a multislice multiple-scattering model to differentiate the emission fluorescence and scattered photons based on the first Born approximation in the incoherent condition^33^ (Fig. 1c and Supplementary Note 1). Then, we applied the alternating direction method of multipliers (ADMM) algorithm^34^ with the multiscale model to update the volumetric fluorescence and 3D scattering potentials iteratively (Supplementary Fig. 2 and Supplementary Video 1).

By modeling both the out-of-focus and scattered photons, we retrieved the 3D fluorescence distribution quantitatively in thick tissue. To show the pipeline, we imaged a GFP-labeled tumor spheroid with a 63 ×/1.4 NA oil-immersion objective at an axial step of 10 μm for 10 subvolumes. Strong background fluorescence was observed in the raw LF images, which was effectively removed by the capability of computational optical sectioning in QLFM (Fig. 1d). Each LF image has a specific axial range for high-resolution 3D components. The large 3D volume with multiple axial steps was reconstructed as a whole during the iterations, making full use of the overlapped axial range and providing a better estimation of the background and uniform resolution at different axial planes (Methods and Supplementary Video 1). In contrast, traditional LFM shows strong artifacts due to crosstalk from the background. The axial side, with larger out-of-focus photons, tends to have higher intensities, demonstrating the loss of the quantitative property in thick tissue.

### Characterization of QLFM

To quantitatively evaluate the improvement of QLFM, we imaged various 2-μm fluorescence beads embedded in a tissue-mimicking phantom made of intralipid and agarose with a 40×/1.0 NA objective and calculated the average SBR at different penetration depths (Methods, Fig. 2a, and Supplementary Fig. 4). By comparing with WFM and different LFM models, shown in Fig. 2b, we found that the naive imaging model of traditional LFM resulted in a similar performance as WFM in tissue penetration. Our multiscale scattering model fully exploits the capability of LFM in deep-tissue imaging with ~20 dB SBR improvement over WFM. Such a great improvement facilitates QLFM high-speed 3D imaging in thick tissue. We then imaged the same densely labeled *Drosophila* brain by QLFM, traditional LFM, WFM, and confocal microscopy for comparison (Supplementary Fig. 5). Despite reduced resolution due to the tradeoff between spatial and angular resolutions in LFM, QLFM showed great sectioning capability comparable to that of confocal microscopy without axial-scanning artifacts, which was much better than those of traditional LFM and WFM.

**Fig. 2.**
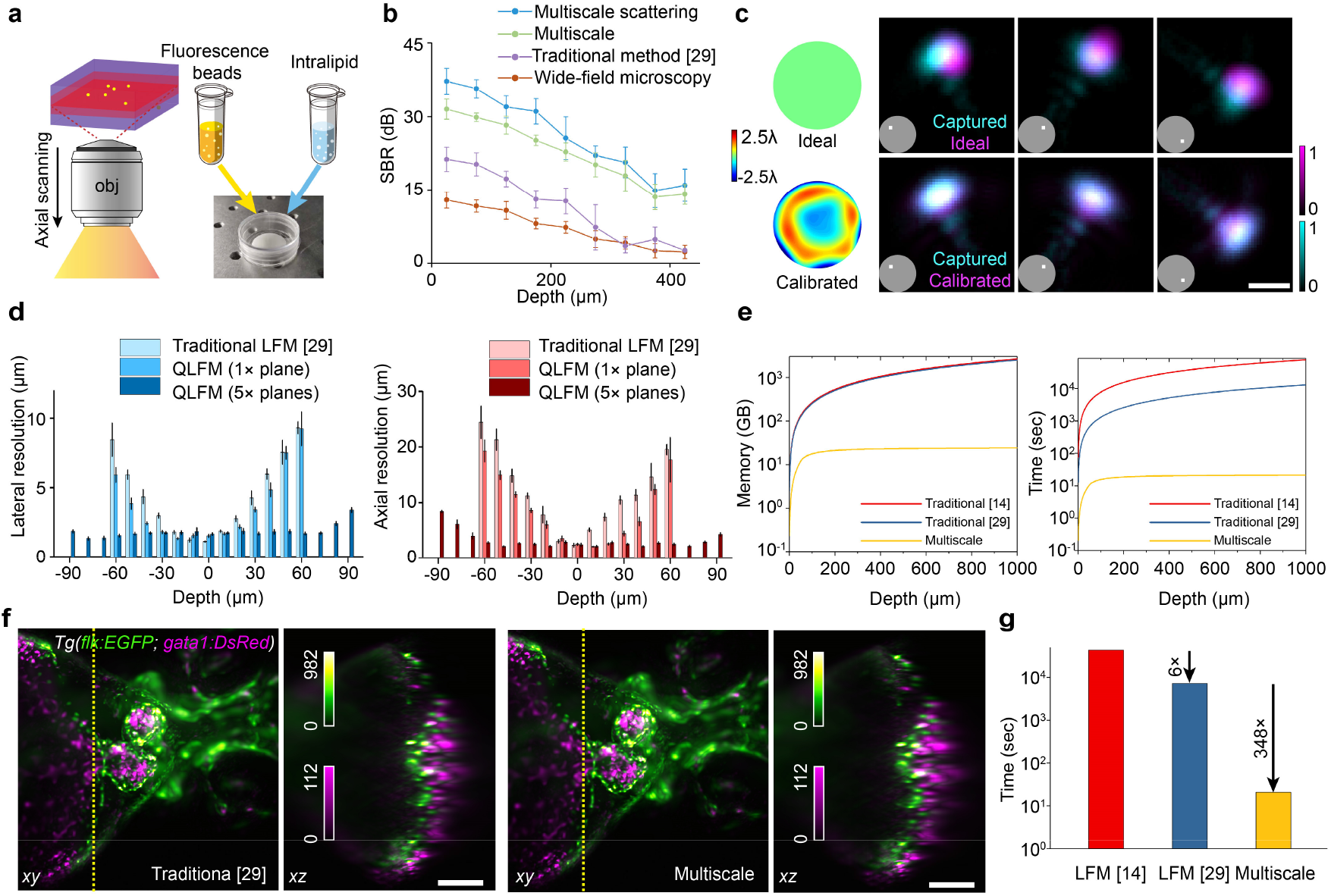
Characterization of quantitative light-field microscopy (QLFM). **a**, Illustrations of the SBR characterization experiment. We fabricated the scattering phantom with the mixture of 0.025% 2-μm fluorescence beads, 1% intralipid, 1% agarose in a petri dish. Then we imaged the sample with a 40×/1.0NA water-immersion objective at different penetration depths with WFM and LFM. **b**, SBR curves of the fluorescence beads at different penetration depths in the intralipid-based tissue-mimicking phantom obtained by WFM, traditional LFM, QLFM with the multiscale model only, and QLFM with the multiscale scattering model. We chose 10 fluorescence beads with the highest fitting degrees for each block covering approximately 40 μm. QLFM shows ~20 dB improvement in the SBR over WFM, indicating an improved penetration depth in deep tissue. **c**, Comparisons among the experimental PSF, the ideal PSF without aberrations, and the calibrated PSF of several selected angular components marked in the inset under the 40×/1.0 NA water-immersion objective. The calibrated wavefront of the system aberration estimated by our iterative phase-retrieval algorithm is shown on the left. **d**, Lateral and axial resolution at different axial planes in traditional LFM and QLFM with 1 and 5 axially scanned planes, which is estimated by the FWHM of subdiffraction-limited fluorescence beads. For each block covering 10 μm, we chose 10 fluorescence beads with the highest fitting degree for statistical analysis. **e**, The curves of the required memory and computation time versus the reconstructed depth range for each volume by different methods with a 20×/0.5NA objective, indicating that the multiscale model is essential to cover a large depth of range for computational optical sectioning. **f**, Orthogonal MIPs of the beating heart in zebrafish larvae reconstructed by different methods with a large axial range. The multiscale model shows similar performance as traditional methods with orders-of-magnitude reduction in computational costs. **g**, The bar graph shows the time required by different methods to reconstruct the volumes shown in f, covering about ~ 700×700×560 μm^3^. Scale bars, 5 μm (**c**) and 100 μm (**f**).

Another problem hindering the quantitative reconstruction in LFM is the inaccurate estimation of the complicated high-dimensional PSF. As shown in Fig. 2c, the system aberration will result in great system errors between the ideal PSF and the experimental PSF measured with a sub-diffraction-limited fluorescence bead. We propose a phase-retrieval-based algorithm to estimate the system aberration by iteratively shrinking the disparity between the simulated PSF and the captured PSF along different angular components with a single image (Supplementary Fig. 6). We find that the calibrated PSF with the aberration wavefront can greatly reduce the reconstruction artifacts close to the native objective plane (Supplementary Fig. 6c). Numerical simulations on different levels of aberrations also shows the effectiveness of the method (Supplementary Fig. 7). We then characterized the system resolution by imaging 500-nm fluorescence beads distributed in 1% agarose with a 40×/1.0 NA water-immersion objective. The average full width at half-maximum (FWHM) was used to estimate the lateral and axial resolutions at different axial planes (Fig. 2d). In addition to the improved resolution, QLFM with high-speed axial scanning at a step size of 30 μm for 5 planes has a uniform resolution of approximately 1.8×1.8×2.5 μm^3^ across a large depth range covering ~ 330×330×180 μm^3^. Moreover, as the multiscale model in phase-space domain avoids the unnecessary up-sampling based on the effective resolution, it can reduce the memory and computing time by orders of magnitude, especially for large depth ranges during reconstruction to get rid of background (Fig. 2e). By modeling the same volume range, our multiscale model with adaptive sampling rate shows the same performance as traditional methods, but reduces the computing time from several hours to several seconds, on a desktop computer equipped with a normal graphical processing unit (CPU: Intel i9-9980XE, RAM: 128 GB, GPU: NVIDIA GeForce RTX 2080 Ti), facilitating the practical use of QLFM.

### High-speed 3D imaging of *Drosophila* embryo and Zebrafish larvae

To demonstrate the superior performance of QLFM over previous methods using compact systems, we performed *in vivo* imaging of various fast biological dynamics in different specimens. First, we imaged the whole-embryo development of histone-labeled *Drosophila* at the millisecond scale with both 40×/1.0 NA (Figs. 3a and b) and 20×/0.5 NA (Figs. 3c and d) objectives for comparison. Different from previous methods capturing only part of the volume at a time^35^, LFM achieves better photon efficiency with low phototoxicity by illuminating and detecting the entire volume simultaneously. While there was almost no contrast in traditional LFM due to the dense labeling and strong scattering, QLFM could still distinguish single cells deep in the embryo at the millisecond scale, extending the applications of LFM to developmental biology.

**Fig. 3.**
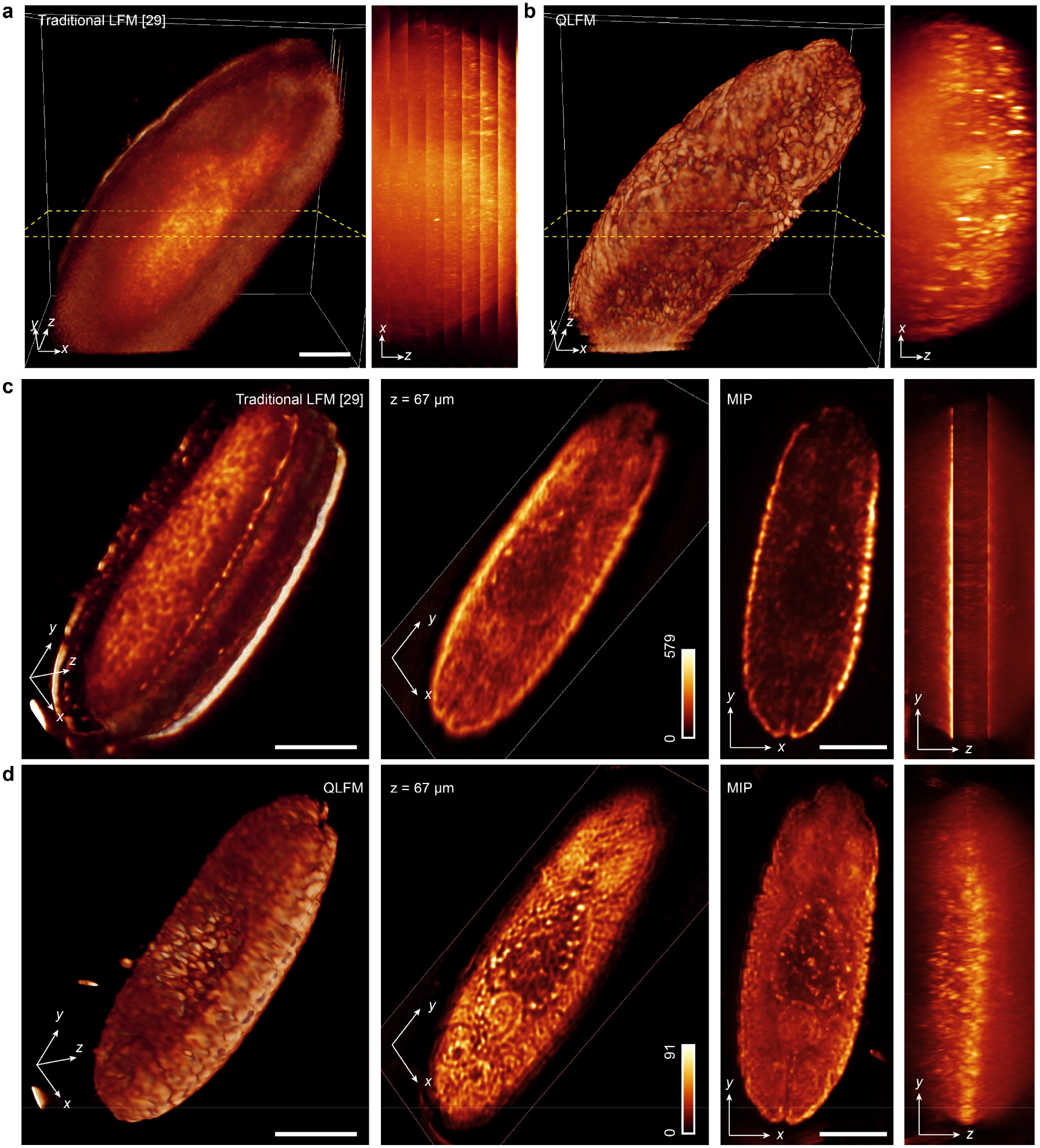
Experimental comparisons on the *Drosophila* embryo with high-speed axial scanning. **a-b**, 3D rendered volumes and 180-μm-xz MIPs at the same time point of traditional LFM and QLFM imaged with a 40×/1.0NA water-immersion objective at a step of 15 μm. QLFM show much better contrast and resolution than traditional LFM without artifacts, illustrating the capability of computational optical sectioning by QLFM with the improved penetration depth. **c-d**, The comparisons between traditional LFM and QLFM with a 20×/0.5NA objective in the form of 3D rendered volumes, slices, and orthogonal MIPs with axial scanning at a step of 50 μm indicating the uniform resolution of QLFM without edge artefacts. Scale bars, 50 μm (**a-b**) and 100 μm (**c-d**).

Second, we performed *in vivo* imaging of heart-beating dynamics in zebrafish larvae, which are difficult to fully capture without simultaneous exposure of the whole volume at high speed. Traditional LFM shows low resolution and contrast with severe artifacts (Fig. 4a), but QLFM, with a more accurate computational model, could obtain artifact-free high-resolution 3D volumes without the requirement of additional light-sheet illumination and multiview objectives (Figs. 4b and c, and Supplementary Figs. 8 and 9). We found successive improvement with increasing model complexity, indicating the necessity of various factors in the model, including out-of-focus fluorescence, sample scattering, and system aberrations (Supplementary Video 2).

**Fig. 4.**
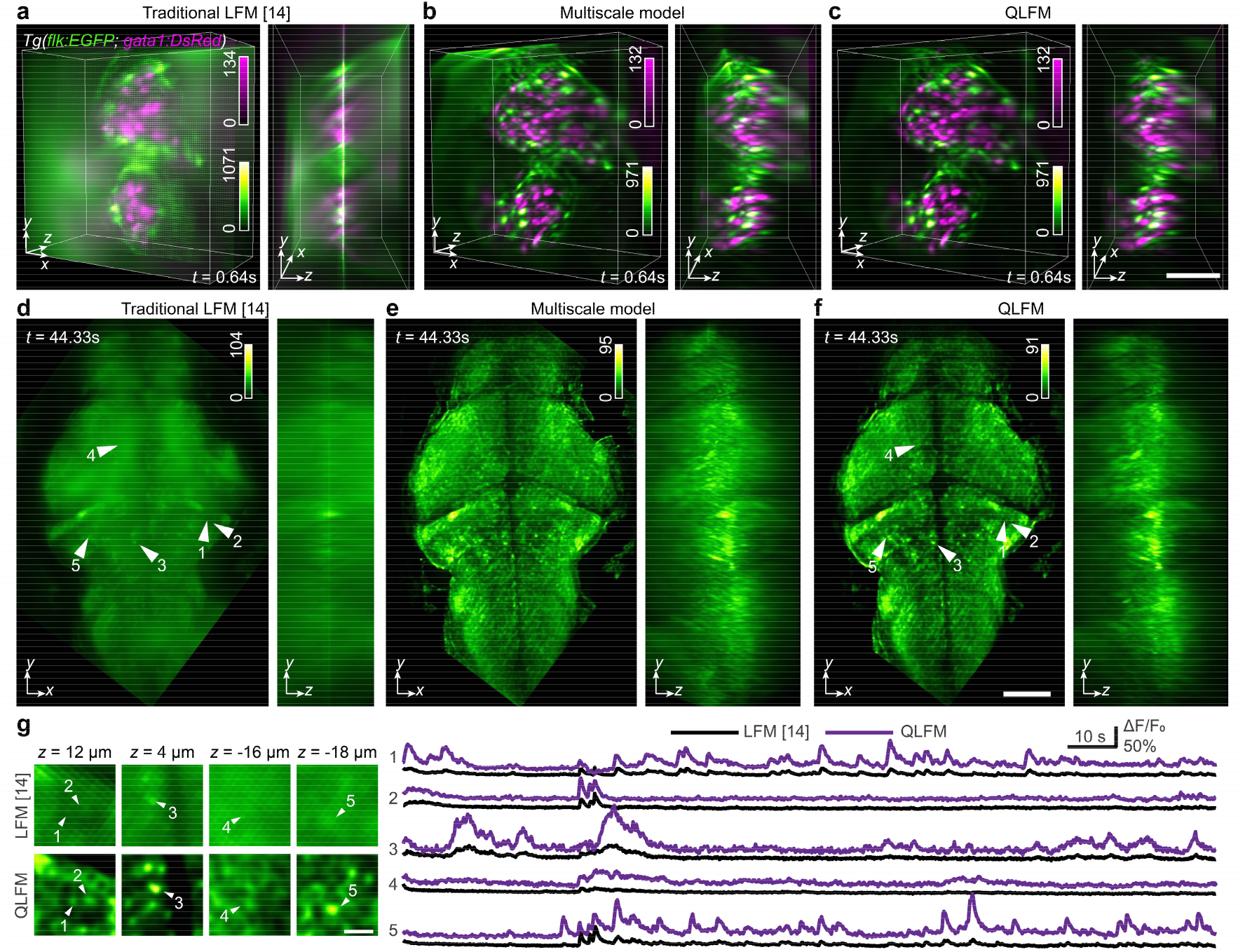
High-speed volumetric imaging in larval zebrafish. **a-c**, 3D rendered volumes of the beating heart at the same time point reconstructed by the traditional LFM, QLFM with the multiscale model only, and QLFM with the multiscale scattering model (Supplementary Video 2). Successive improvements can be observed with significantly reduced artifacts and background for quantitative imaging. **d-f**, Orthogonal maximum-intensity projections (MIP) of the whole brain at the same time point reconstructed by the traditional LFM, QLFM with the multiscale model only, and QLFM with the multiscale scattering model, illustrating the effectiveness of computational optical sectioning in densely labeled samples (Supplementary Video 3). **g**, Zoom-in slices of the same neurons marked in d and f and their temporal traces for comparison, illustrating the improved SBR. Scale bars, 50 μm (**a-c**), 100 μm (**d-f**) and 20 μm (**g**).

Whole-brain neural recording in zebrafish larvae is another typical application of LFM, but the imaging performance is far from satisfactory due to the densely packed neurons without complicated systems^14^ (Fig. 4d). Although the spatiotemporal sparsity facilitates the accurate localization and extraction of neural signals^36,37^, it is still susceptible to nonuniform background fluorescence and inevitable structural changes in living animals. By imaging the whole-brain neural activities in zebrafish larvae at 24 Hz with a 20×/0.5 NA objective, we demonstrate that QLFM can achieve effective single-neuron resolution with proper models to remove both out-of-focus and scattered photons for each single frame without the requirement to calculate the standard deviation of many frames for better contrast (Figs. 4e and f, and Supplementary Video 3). The temporal traces of several neurons with simple region-of-interest (ROI) averaging demonstrate the increased contrast and reduced crosstalk between adjacent neurons in QLFM (Fig. 4g).

### Large-scale 3D calcium imaging in mice brain

The mammalian brain is a more challenging case due to its strong scattering and dense neural structures. We tested the performance of QLFM by imaging the 3D calcium activities in an awake head-fixed mouse, which was labeled with GCaMP6s by virus injection, under a 20×/0.5 NA objective. To visualize the 3D distribution of neurons, traditional LFM usually requires the standard deviation of thousands of frames to reduce the background (Supplementary Fig. 10a). However, the 3D signals of a single frame are usually flooded in the background fluctuations due to the low SBR (Fig. 5a). In contrast, with the capability of computational optical sectioning, QLFM shows significantly improved contrast (Fig. 5b and Supplementary Video 4). In addition to the detection of more neurons (Supplementary Fig. 10a), QLFM allows the measurement of calcium dynamics in a quantitative manner without the need for additional background subtractions or filtering to improve the contrast, making it more reliable for large-scale neural recording (Fig. 5c). Background fluorescence usually increases with the increasing of NA due to the smaller depth of field, especially for the wide-field imaging. We therefore imaged the same mouse with a 40×/1.0 NA water-immersion objective at different depths (Supplementary Figs. 10b-e and Supplementary Video 5). QLFM not only resolved more neurons with better resolution and contrast than traditional LFM but also showed stable performance at different depths up to 300 μm (Figs. 5d and e).

**Fig. 5.**
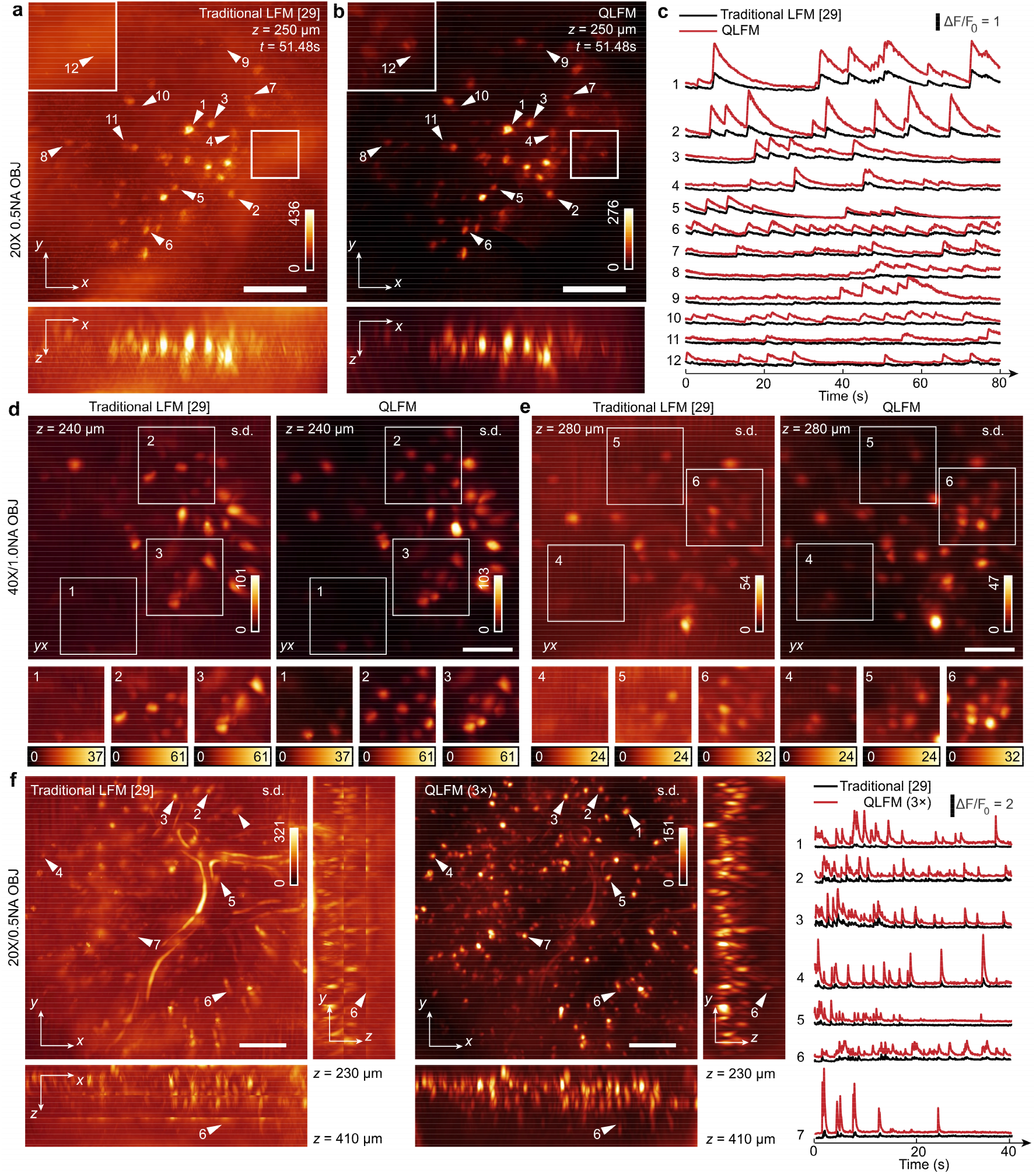
3D functional imaging in awake mouse brains. **a-b**, Orthogonal MIPs of GCaMP6s-labeled neurons at *t*=51.48 s reconstructed by traditional LFM and QLFM with a 20×/0.5NA objective, indicating the reduced background with computational optical sectioning (Supplementary Video 4). More neurons were revealed by QLFM in the mouse cortex with the native objective plane at a depth of ~ 250 μm. **c**, Temporal traces of the marked neurons in a and b, demonstrating the quantitative calcium responses in QLFM with improved contrast. **d-e**, Orthogonal MIPs of the standard deviation across 2000 volumes imaged at a center depth of240 μm and 280 μm in the cortex, respectively, under a 40×/1.0 NA water-immersion objective (Supplementary Video 5). **f**, Orthogonal MIPs of the standard deviation across 500 volumes for GCaMP6f-labeled L2/3 neurons in an awake behaving mice. The video was captured with high-speed axial scanning at a step of 50 μm for 3 planes at 24 Hz with a 20×/0.5 NA objective. Temporal traces of several marked neurons at different depths are shown on the right. QLFM shows uniform resolution and consistent calcium responses across a large depth range, while there is barely contrast in traditional LFM due to severe background. Scale bars, 100 μm (**a-b, f**) and 50 μm (**d-e**).

By high-speed axial scanning, we could further increase the axial coverage with redundant 3D imaging speed for calcium dynamics. We then conducted millisecond-scale calcium imaging in awake double-transgenic Rasgrf2-2A-dCre/Ai148D mice across a large volume covering ~ 700×700×180 μm^3^ by high-speed scanning of 3 planes. While there was barely contrast in traditional LFM due to the severe artifacts especially in the axial domain, which was similar to previous work^36^, QLFM showed a much larger penetration depth with uniform resolution (Figs. 5f and g and Supplementary Video 6). Even the neurons located ~400 μm deep exhibited remarkable calcium responses.

## Discussion

In summary, we developed an incoherent multiscale scattering model to fully exploit the intrinsic high-dimensional property of phase-space measurements and achieved computational optical sectioning, facilitating high-speed, large-scale, quantitative 3D imaging in deep tissue with a compact system.

Different from wide-field imaging with a tight focal plane, phase-space imaging provides a tomographic framework by keeping the photons focused along different angles with an extended depth of field. Such a process makes the signals originated from different axial planes show much more distinguishable features than WFM, which can be utilized to remove background fluorescence far from the native objective plane computationally, as long as we model the imaging process in a complete space. Additional confocal rejection or light-sheet illuminations will definitely further reduce the shot-noise fluctuations by getting rid of the background photons physically but at the cost of system compactness, space constraints, or detection parallelization.

In addition, as a novel computational model, our method is compatible with different phase-space imaging schemes without the requirement of any hardware modifications. In combination with the low phototoxicity and high photon efficiency, the orders-of-magnitude reductions in background fluorescence, artifacts, and computational costs enable the practical and versatile applications of QLFM as a compact and robust add-on to normal wide-field microscopy, making the advanced imaging capability generally accessible to the broad biology community.

## Methods

### Experimental setup

Our QLFM system works as an add-on to a normal epifluorescence microscope, with a microlens array inserted at the image plane. A customized upright microscope was used for mouse experiments, while a commercial inverted microscope (Zeiss Observer Z1) was used for the others. Another 4f system (with a magnification of 0.845) relayed the back focal plane of the microlens array (with a pitch size of 100 μm and focal length of 2.1 mm) to the camera (Andor Zyla 4.2 plus, 2,048 × 2,048 pixels) so that each microlens covered ~13 × 13 sensor pixels, corresponding to a 2.15 μm × 2.15 μm area at the sample plane for the 40×/1.0 NA water-immersion objective. Multiple lasers (488 nm and 561 nm) were used for the fluorescence excitation of multiple channels, which were synchronized with the camera for time division multiplexing. A piezo objective scanner (PI P-725.4CD) was used for high-speed axial scanning with a resolution of 1.25 nm at 100 Hz. Detailed imaging conditions and reconstruction parameters for all fluorescence experiments in the paper, including the excitation power, exposure time, frame rate, voxel size, fluorophore, protein, filter set, and objective, are illustrated in Supplementary Table 1.

### 3D deconvolution with a multiscale scattering model

To reconstruct the 3D fluorescence information in deep tissue in a quantitative way, we propose a novel 3D deconvolution algorithm with a multiscale scattering model to iteratively update both the emission fluorescence and scattering photons across an extremely large depth range (Fig. 1c). The whole pipeline of the algorithm with the pseudocode is shown in Supplementary Fig. 2.

With simple pixel realignment, the raw LF data can be represented as multiple angular components, or smoothed phase-space measurements^20^, which can be used for phase-space deconvolution^29^ to reduce artifacts and increase the convergence speed. However, the reconstructed volume, regardless of the deconvolution algorithm^29,30^, is limited to only dozens of axial planes due to the heavy computational cost. Therefore, the out-of-focus fluorescence has usually been modeled as a constant in previous methods, which is fine for thin samples but leads to severe artifacts and background noise in deep tissue with nonuniform out-of-focus distributions. Fortunately, both the lateral and axial resolution of LFM will gradually decrease with increasing distances from the native objective plane. We can establish a multiscale grid to sample a large 3D volume at different intervals based on the characterized resolution with an exponential fit (Supplementary Fig. 2). The out-of-focus fluorescence with a nonuniform 3D distribution can then be estimated with the depth-dependent features within a large depth of field (Supplementary Fig. 1), akin to structured illumination microscopy exhibiting computational optical sectioning. As this method does not require dense axial sampling and a lateral sampling rate as high as the camera pixel number for each plane, which was necessary in previously reported methods due to the spatially variant PSF^30^, the unknown variables can be reduced by orders of magnitude to accelerate the reconstruction speed when modeling the out-of-focus fluorescence of a much larger volume (Fig. 2e). For the same volume size, our multiscale model can achieve almost the same performance as traditional dense-sampling models at a much faster speed. In contrast, the traditional method^29^, without a sufficient axial range in the model, shows apparent artifacts and an increase in the background.

Scattering is a recurring challenge in thick tissue and is the fundamental limitation of the penetration depth in light microscopy, leading to reduced resolution and contrast^38^. Although the multilayer scattering model has recently achieved great success in coherent imaging modalities, such as diffraction tomography^39^, the spatially nonuniform influence of scattering is barely considered in fluorescence imaging. LFM provides a great opportunity to differentiate emission fluorescence and scattered photons with its 4D phase-space measurements. We propose a multilayer scattering model based on the first Born approximation in incoherent conditions by describing the relation between the emission fluorescence *I*^(*i*)^(**r**) and scattered photons *I*^(*s*)^(**r**) as follows:

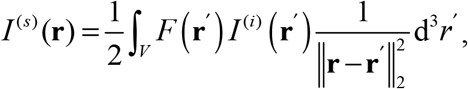

where *F* is defined as the scattering potential energy at the 3D position *r*, and *V* is the whole 3D volume range. The detailed derivations and discretized version are illustrated in Supplementary Note 1. Then, an ADMM framework^34^ is used to iteratively update the sample information and the scattered photons to retrieve the quantitative fluorescence distributions in deep tissue with increased contrast and resolution (Supplementary Fig. 2). The reconstruction time of a single volume with a full FOV of the sensor used here is about several seconds for a normal desktop computer with a graphical processing unit.

### 3D deconvolution with axial scanning for LFM

The high-resolution 3D range of LFM for a snapshot is fixed for a specific objective. High-speed axial scanning at a large step size is a straightforward method to flexibly adjust the depth range and 3D imaging speed. Such a capability is important for versatile applications with different requirements for the volume size and imaging speed. However, traditional methods show severe artifacts at the stitching edges at low contrast (Figs. 1d and 5f), as they usually reconstruct each subvolume based on every single LF image separately and stitch the subvolumes in the axial domain^36^. Here, with the multiscale model in QLFM, we take the entire 3D volume as a whole with the multiscale sampling rate during reconstruction. As shown in Supplementary Video 1, after pixel realignment of all the raw LF images, we can achieve multiple focal stacks for different angular components, making full use of the intrinsic continuous property of different angular components in the axial domain. Due to the large depth of field for each angular component in QLFM, we can use a large step for axial scanning with much less time required to cover the same volume than in WFM. Then we update the entire large volume as a whole along different angles, viewing each angular focal stack as the minimum unit to calculate the error map for each iteration. Such a process is akin to wide-field deconvolution first for each angle followed by tomographic reconstruction later for different angles. Finally, the same ADMM framework is applied to the entire volume to update the emission fluorescence and scattered photons iteratively. In this case, uniform resolution can be achieved at different axial planes across a large depth range without any artifacts (Fig. 1d and 2d). In addition, the computational cost can be further reduced, as we can conduct 3D deconvolution with the phase-space PSF for each angular focal stack.

### Phase-retrieval-based PSF calibration

The PSF of traditional LFM is calculated based on wave optics theory in an ideal imaging condition^29,30^. However, the experimental imaging system usually has complicated system aberrations, which will not only reduce the imaging resolution but also introduce severe reconstruction artifacts close to the native objective plane. Here, we propose a phase-retrieval-based algorithm to calibrate the experimental PSF with a single image of subdiffraction-limited fluorescence beads (Supplementary Fig. 6). The simulated PSF is first initialized with experimental parameters without any aberration wavefront. Then, we calculate the correlations between the captured image and the simulated PSF along different angular components to estimate the wavefront error at different sub-apertures. The aberrated wavefront of the whole NA is then integrated from the correlation map for continuous distributions. The estimated wavefront is fitted with Zernike polynomials and fed into the wave optics model to generate the new simulated PSF, which is used again to match with the captured PSF. The calibrated PSF is iteratively updated through the above process until the phase map converges, which usually takes approximately 3~4 iterations. Interestingly, we found that the experimental system with spherical aberrations can remove the reconstruction artifacts close to the native objective plane as long as we had an accurate estimation of the aberration wavefront.

### Fluorescence bead preparation for system characterization

For resolution characterization, 500-nm yellow-green fluorescent microspheres (Thermo Scientific, FluoSpheres, carboxylate-modified microspheres) were mixed with 1% agarose at a ratio of 1:1,000,000. We placed the mixture in a 35-mm petri dish and captured ~100 LF images for statistical analysis with a 40×/1.0 NA water-immersion objective. We then calculated the FWHM of the lateral and axial profiles of the reconstructed beads on different axial planes. For each block covering 10 μm, we chose 10 beads with the highest fitting degree to calculate the mean and standard deviation.

For SBR characterization, we fabricated a scattering phantom with the mixture of 1% agarose, 1% intralipid (Absin 68890-65-3, 20% emulsion), and 0.025% 2-μm fluorescence beads, which was placed in a 35-mm petri dish. The 2-μm fluorescence beads (Thermo Scientific, FluoSpheres, carboxylate-modified microspheres) were randomly distributed in the intralipid and imaged using a 40×/1.0 NA water-immersion objective. The reconstructed mean intensity of the beads was viewed as the signal, while the mean intensity of the reconstructed background without samples was viewed as the background. Several reconstructed slices at different imaging depths are shown in Supplementary Fig. 4.

### Tumor spheroid preparation

B16 cells (ATCC^®^ CRL-6475™, mouse skin melanoma cells) were purchased from ATCC and cultured in RPMI 1640 medium supplemented with 10% FBS, 1% pen/strep antibiotics and 1% NEAA (all from GIBCO). Cells were then transfected with the EGFP-PH plasmid (Addgene Plasmid #96948), and stable EGFP-expressing B16 cells were selected by neomycin (G418) and maintained. To prepare tumor spheroids, 4 × 10^3^ EGFP-expressing B16 cells per well were seeded in round-bottomed 96-well plates (Corning) and cultured in RPMI 1640 medium supplemented with 10% FBS, 2% B-27 supplement (GIBCO), 2% methyl cellulose (Sigma-Aldrich), 1% pen/strep antibiotics and 1% NEAA. After 2 days, each formed spheroid was transferred as 1 spheroid per well and cultured for another 2 days. During LFM imaging, the GFP-B16 tumor spheroids were transferred to Lab-Tek II cover-glass-bottomed 8-chamber slides and imaged in HBSS supplemented with 2% FBS (all from Invitrogen) using a 63×/1.4 NA oil-immersion objective.

### *In vivo* imaging of zebrafish larvae

All zebrafish experimental procedures were conducted with ethical approval from the Animal Care and Use Committee of Tsinghua University. For imaging of the vasculature and blood flow dynamics, *Tg(flk:EGFP; gata1: DsRed)* transgenic zebrafish embryos were collected and cultured at 28.5 °C in Holtfreter’s solution. At 4–5 days postfertilization (dpf), the zebrafish larvae were anesthetized by ethyl 3-aminobenzoate methanesulfonate salt (100 mg/L) and mounted in 1% low-melting-point agarose for imaging at 26–27°C. For whole-brain calcium imaging, *Tg(huc:GCaMP6)* transgenic zebrafish embryos were collected and kept at 28.5 °C in Holtfreter’s solution. At 4-5 dpf, the larvae were mounted in 1% low-melting-point agarose for imaging at 26–27 °C.

### Preparation of fixed *Drosophila* brain samples

All Drosophila experimental procedures were conducted with ethical approval from the Animal Care and Use Committee of Tsinghua University. Female Drosophila brains were dissected and fixed in 4% paraformaldehyde (PF A, Cat# AR-0211, Dingguo Biotech, China) at room temperature for 30 mins on a shaker. Each brain was rehydrated with 0.3% Triton X-100 (Solarbio 524A0513) in phosphate-buffered saline (PBS) for 4×20 mins at room temperature and then incubated in block solution (5% goat serum in washing buffer) for 30 mins at room temperature. The brain was then incubated overnight with primary antibody (mouse monoclonal nc82 (Developmental Studies Hybridoma Bank)), which was diluted at 1:500 in block solution at 4 °C. The brain was then washed in 0.5% PBST for 3×1 hour at room temperature. Finally, the brain was mounted directly for imaging by LFM.

### Imaging of *Drosophila* embryos

All *Drosophila* embryos (Fig. 3) expressed histones tagged with EGFP (His2Av, BL24163). The collection and preparation of *Drosophila* embryos were performed according to the commonly used protocol^40^. Two-hour *Drosophila* embryos were collected within a specific collection chamber. After incubation at 25 °C for 10 hours, each embryo was attached to a glass microscope slide with double-sided sticky tape. We used forceps to carefully roll the embryo on the tape until its chorion popped open. Then, the embryo was transferred to the glue line on a dish (Ibidi μ-Dish, 35 mm, high) and covered with mineral oil (Sigma-Aldrich, Halocarbon 27 oil) for live imaging.

### Mouse experiments

All procedures involving mice were approved by the Institutional Animal Care and Use Committee of Tsinghua University. We used both male and female C57BL/6 mice 10 weeks to 6 months old without randomization or blinding. We performed the craniotomy as previously described^41^, with a window size of ~8 mm×8 mm. Then, we installed a flat optical window and cemented a custom-made coverslip (D-shape) and aluminum head posts to the skull. For acute imaging, we used adult double-transgenic Rasgrf2-2A-dCre/Ai148D mice (JAX No.: 022864 and 030328) to specifically label cortical layer 2/3 neurons^42^. For chronic imaging, adult C57BL/6 mice injected with diluted AAV9-hSyn-GCaMP6s virus (from BrainVTA Technology Co., Ltd., China) were allowed to recover for at least 2 weeks after craniotomy. During LFM imaging, awake mice were placed in a tube with the head restrained under the objective.

### Data analysis

All data analyses were performed with customized MATLAB (MathWorks, MATLAB 2018) programs and Amira (Thermo Fisher Scientific, Amira 2019). The hardware was controlled using LabVIEW 2018 (National Instruments). The 3D rendering of the volumes in figures and videos was performed by Amira. The 3D tracking of 7 representative blood cells in the heart of the zebrafish larvae was carried out manually in MATLAB.

### Neural activity extraction

The calcium responses in both zebrafish and mice were extracted directly through signal averaging of the manually selected ROIs covering the selected neurons. The ROIs were ~8×8×10 μm^3^ in size for zebrafish larvae and ~10×10×10 μm^3^ in size for mice to match the size of the neuron. The temporal traces of neural activity were calculated by *ΔF/F_0_*= (*F-F_0_*)/*F_0_*, where *F* is the raw averaged intensity of the ROI, and *F_0_* is the baseline fluorescence intensity. To estimate *F_0_* for each ROI, we first calculated the average intensity of the trace and averaged all time points with signals below 120% of the calculated average.

### Data availability

All relevant data are available from the corresponding authors upon reasonable request.

### Code availability

We will open all of codes with example datasets in Google Drive and Github after the paper is accepted.

## Acknowledgments

We thank Li Yu, Jiesi Feng, Yulong Li, Jing He, Guihua Xiao, and Hao Xie for their assistance in sample preparation. This work was supported by the National Natural Science Foundation of China (61327902, 61827804, 61620106005), the Beijing Municipal Science & Technology Commission (BMSTC) (No. Z181100003118014), the National Key Research and Development Program of China (2020AAA0130000), the Postdoctoral Science Foundation of China (2019M660644), and the Tsinghua University Initiative Scientific Research Program. J.W. sincerely appreciates funding from the National Postdoctoral Program for Innovative Talent.

## Author contributions

Q.D., X.J., and J.W. conceived and designed the project. Q.D. and X.J. supervised the research. Z.L. and J.W. designed and built the optical system. Y.Z., X.J. and J.W. conducted the numerical simulation and developed the deconvolution algorithm with the multiscale scattering model. J.W. and D.J. designed the biological experiments. X.L. conducted the hardware synchronization. J.X. derived the scattering model in incoherent conditions. T.Z. designed the graphical user interface for software control. D.J. prepared the zebrafish larvae. Z.L., Y.Z., D.J. and J.W. conducted most biological experiments, data collection and volume reconstructions. X.L. and Y.C. conducted algorithm performance optimization. Y.Z. designed the system calibration method. J.W., Z.L., Y.Z., X.J., and Q.D. prepared figures and wrote the manuscript with input from all authors.

## Competing financial interests

The authors declare no competing financial interests.

## Materials & correspondence

Correspondence and requests for materials should be addressed to (Q.D.), (X.J.), and (J.W.).

## Notes

### Competing Interest Statement

The authors have declared no competing interest.

## References

1. Weisenburger, S. & Vaziri, A. A guide to emerging technologies for large-scale and whole-brain optical imaging of neuronal activity. Annu. Re. Neurosci. 41, 431–452 (2018).

2. Ji, N., Freeman, J. & Smith, S. L. Technologies for imaging neural activity in large volumes. Nat. Neurosci. 19, 1154 (2016).

3. Pittet, M. J. & Weissleder, R. Intravital imaging. Cell 147, 983–991 (2011).

4. Pawley, J.B. Handbook of Biological Confocal Microscopy, edn 2. (Plenum Press, New York, 1995).

5. Shimozawa, T. et al. Improving spinning disk confocal microscopy by preventing pinhole cross-talk for intravital imaging. Proc. Natl. Acad. Sci. U. S. A. 110, 3399–3404 (2013).

6. Helmchen, F. & Denk, W. Deep tissue two-photon microscopy. Nat. Methods 2, 932–940 (2005).

7. Zhu, G., van Howe, J., Durst, M., Zipfel, W. R. & Xu, C. Simultaneous spatial and temporal focusing of femtosecond pulses. Opt. Express 13, 2153–2159 (2005).

8. Bouchard, M. B. et al. Swept confocally-aligned planar excitation (SCAPE) microscopy for high-speed volumetric imaging of behaving organisms. Nat. Photonics 9, 113–119 (2015).

9. Ahrens, M. B., Orger, M. B., Robson, D. N., Li, J. M. & Keller, P. J. Whole-brain functional imaging at cellular resolution using light-sheet microscopy. Nat. Methods 10, 413–420 (2013).

10. Wu, Y. & Shroff, H. Faster, sharper, and deeper: structured illumination microscopy for biological imaging. Nat. Methods 15, 1011–1019 (2018).

11. Mertz, J. Optical sectioning microscopy with planar or structured illumination. Nat. Methods 8, 811–819 (2011).

12. Winter, P.W. & Shroff, H. Faster fluorescence microscopy: advances in high speed biological imaging. Curr. Opin. Chem. Biol. 20, 46–53 (2014).

13. Laissue, P. P. et al. Assessing phototoxicity in live fluorescence imaging. Nat. Methods 14, 657–661 (2017).

14. Prevedel, R. et al. Simultaneous whole-animal 3D imaging of neuronal activity using light-field microscopy. Nat. Methods 11, 727–730 (2014).

15. Alonso, M. Wigner functions in optics: describing beams as ray bundles and pulses as particle ensembles. Adv. Opt. Photon. 3, 272–365 (2011).

16. Waller, L., Situ, G. & Fleischer, J. W. Phase-space measurement and coherence synthesis of optical beams. Nat. Photon. 6, 474–479 (2012).

17. Liu HY, Jonas E, Tian L, Zhong JS, Recht B et al. 3D imaging in volumetric scattering media using phase-space measurements. Opt. Express 2015; 23: 14461–14471.

18. Takasaki, K. T. & Fleischer, J. W. Phase-space measurement for depth-resolved memory-effect imaging. Opt. Express 22, 31426–31433 (2014).

19. Levoy, M., Ng, R., Adams, A., Footer, M. & Horowitz, M. Light field microscopy. ACM Trans. Grapgh. 25, 924–934 (2006).

20. Tian, L., Zhang, Z., Petruccelli, J. C. & Barbastathis, G. Wigner function measurement using a lenslet array. Opt. Express 21, 10511–10525 (2013).

21. Wagner, N. et al. Instantaneous isotropic volumetric imaging of fast biological processes. Nat. Methods 16, 497–500 (2019).

22. Lin, Q. et al. Cerebellar Neurodynamics Predict Decision Timing and Outcome on the Single-Trial Level. Cell 180, 536–551 (2020).

23. Abrahamsson, S. et al. Fast multicolor 3D imaging using aberration-corrected multifocus microscopy. Nat. Methods 10, 60–63 (2013).

24. Yang, W. et al. Simultaneous Multi-plane Imaging of Neural Circuits. Neuron 89, 269–284 (2016).

25. Xiao, S. et al. High-contrast multifocus microscopy with a single camera and z-splitter prism. Optica 7, 1477–1486 (2020).

26. Taylor, M. A., Nöbauer, T., Pernia-Andrade, A., Schlumm, F. & Vaziri, A. Brain-wide 3D light-field imaging of neuronal activity with speckle-enhanced resolution. Optica 5, 343–353 (2018)

27. Wagner, N. et al. Instantaneous isotropic volumetric imaging of fast biological processes. Nat. Methods 16, 497–500 (2019).

28. Zhang, Z. et al. Imaging volumetric dynamics at high speed in mouse and zebrafish brain with confocal light field microscopy. Nat. Biotechnol. 1–10 (2020).

29. Lu, Z. et al. Phase-space deconvolution for light field microscopy. Opt. Express 27, 18131–18145 (2019).

30. Broxton, M. et al. Wave optics theory and 3-D deconvolution for the light field microscope. Opt. Express 21, 25418–25439 (2013).

31. Gustafsson, M. G. L. Nonlinear structured-illumination microscopy: Wide-field fluorescence imaging with theoretically unlimited resolution. Proc. Natl. Acad. Sci. U.S.A. 102, 13081–13086 (2005).

32. Tian, L. & Waller, L. 3D intensity and phase imaging from light field measurements in an LED array microscope. Optica 2, 104–111 (2015).

33. Born, M., Wolf, E. Principles of Optics: Electromagnetic Theory of Propagation, Interference and Diffraction of Light. Elsevier (2013).

34. Boyd, S., Parikh, N., Chu, E., Peleato, B. & Eckstein, J. Distributed optimization and statistical learning via the alternating direction method of multipliers. Foundations and Trends in Machine Learning 3, 1–122 (2010).

35. Royer, L. A. et al. Adaptive light-sheet microscopy for long-term, high-resolution imaging in living organisms. Nat. Biotechnol. 34, 1267–1278 (2016).

36. Nöbauer, T. et al. Video rate volumetric Ca2+ imaging across cortex using seeded iterative demixing (SID) microscopy. Nat. Methods 14, 811 (2017).

37. Pégard, N. C. et al. Compressive light-field microscopy for 3D neural activity recording. Optica 3, 517–524 (2016).

38. Theer, P. & Denk, W. On the fundamental imaging-depth limit in two-photon microscopy. J. Opt. Soc. Am. A 23, 3139–3149 (2006).

39. Chen, M., Ren, D., Liu, H.-Y., Chowdhury, S. & Waller, L. Multi-layer Born multiple-scattering model for 3D phase microscopy. Optica 7, 394–403 (2020).

40. Conduit, P. T., Hayward, D. & Wakefield, J. G. Microinjection techniques for studying centrosome function in Drosophila melanogaster syncytial embryos. Methods Cell Biol. 129, 229–249 (2015).

41. Chen, T. W. et al. Ultrasensitive fluorescent proteins for imaging neuronal activity. Nature 499, 295–300 (2013).

42. Daigle, T. L. et al. A Suite of Transgenic Driver and Reporter Mouse Lines with Enhanced Brain-Cell-Type Targeting and Functionality. Cell 174, 465–480 (2018).

