## supplementary information for "Computational optical sectioning by phase-space imaging with an incoherent multiscale scattering model"

| | |
|---|---|
| **Supplementary Figure 1** | Illustrations of the extended depth of field (DOF) by phase-space imaging for background modeling |
| **Supplementary Figure 2** | The pseudo codes (left) and corresponding illustrations (right) of 3D deconvolution with a multiscale scattering model |
| **Supplementary Figure 3** | Numerical analysis of the quantitative property of QLFM and traditional LFM with increasing background levels |
| **Supplementary Figure 4** | Characterization of the signal-to-background ratio (SBR) in the tissue-mimicking phantom |
| **Supplementary Figure 5** | Experimental comparisons among confocal microscopy, WFM, traditional LFM, and QLFM on the same fixed *Drosophila* brain |
| **Supplementary Figure 6** | The pseudo codes and corresponding illustrations for phase-retrieval-based PSF calibration |
| **Supplementary Figure 7** | Quantitative analysis of the phase-retrieval-based PSF calibration algorithm |
| **Supplementary Figure 8** | Experimental comparisons on the beating heart in zebrafish larvae to show successive improvements |
| **Supplementary Figure 9** | Experimental comparisons on the large-scale blood flow in zebrafish larvae between traditional LFM and QLFM |
| **Supplementary Figure 10** | *In vivo* 3D calcium imaging in mouse cortex |
| **Supplementary Note 1** | Incoherent scattering model for 3D deconvolution |
| **Supplementary Table 1** | Imaging conditions for all fluorescence experiments |
| **Supplementary Video 1** | Concept and pipeline of the multiscale scattering model for 3D deconvolution |
| **Supplementary Video 2** | Heart-beating dynamics in zebrafish larvae imaged at 25 Hz by two channels to show the successive improvements with the model complexity |

| | |
|---|---|
| **Supplementary Video 3** | Experimental comparisons on whole-brain calcium imaging of zebrafish larvae at 24Hz |
| **Supplementary Video 4** | Experimental comparisons on 3D calcium imaging of a virus-injected awake mouse with a 20×/0.5 NA objective |
| **Supplementary Video 5** | Experimental comparisons on 3D calcium imaging of a virus-injected awake mouse with a 40×/1.0 NA water-immersion objective |
| **Supplementary Video 6** | Experimental comparisons on 3D calcium imaging of a transgenic awake mouse with high-speed axial scanning |

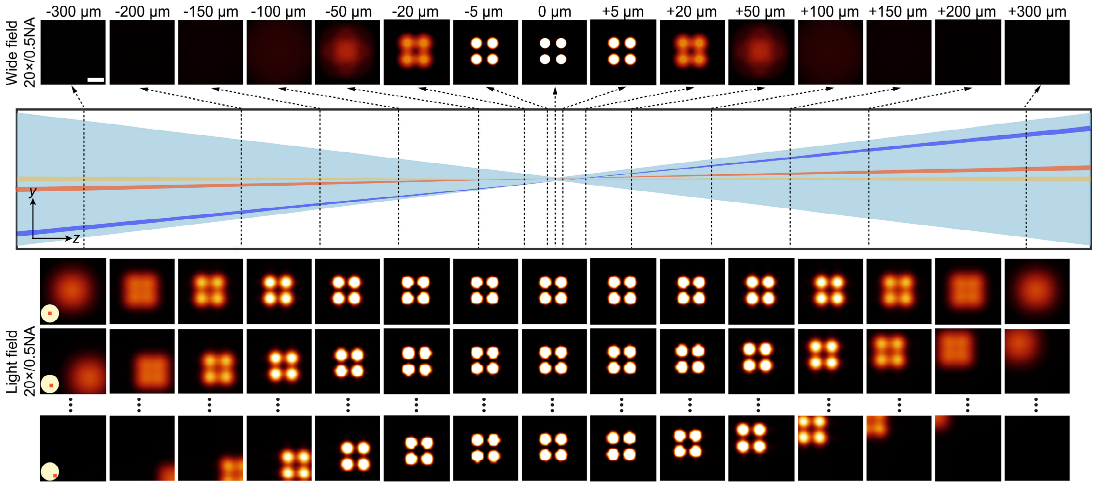


**Supplementary Figure 1 | Illustrations of the extended depth of field (DOF) by phase-space imaging for background modeling.** We show the simulated images of four 8-μm fluorescence beads placed at different axial positions imaged with a 20×/0.5NA objective by wide-field microscopy (WFM) and light-field microscopy (LFM), respectively. The LF images of specific angular components after pixel realignment are shown for comparisons. The photons disperse quickly in WFM with the increase of the out-of-focus distances, generating a uniform background, which is hard to be removed without contrast. On the contrary, the photons in LFM are kept focused within an extended DOF and have unique depth-dependent shifts in different angular components, which can be easily rejected once modeled in the algorithm. Scale bar, 20 μm.

**Algorithm** 3D deconvolution with multiscale scattering model

**Input:** point spread function $H(\rho,u,nz)$, axial downsample rate $r_z$, lateral downsample rate $r_{xy}$, Spatial-angular components $W(\rho,u)$, maximum iteration $M$, initial volume $X_{all}(\rho,nz)$, initial scattering volume $I^{(s)}(\rho,nz)$, Green function $G$, update rate $\lambda$, ratio $\beta$, learning rate $\alpha$

**Output:** descattered 3D volume $X(\rho,z)$

```
H(ρ,u,nz) = DS_xy(DS_z(H(ρ,u,nz),r_z),r_xy)
X_all(ρ,nz) = DS_xy(DS_z(X_all(ρ,nz),r_z),r_xy)
I^(s)(ρ,nz) = DS_xy(DS_z(I^(s)(ρ,nz),r_z),r_xy)

for m = 1:M do
    for u = 1:size(H,3) do
        if (m==1 && u ==1) do
            X(ρ,nz) = RL_deconv(H_u(ρ,nz), W_u(ρ), X(ρ,nz), λ)
                // Update X(ρ,nz) along u with RL deconvolution
            X_all(ρ,nz) = X(ρ,nz)
        end if
        X_all(ρ,nz) = RL_deconv(H_u(ρ,nz), W_u(ρ), X_all(ρ,nz), λ)
            // Update X_all(ρ,nz) along u with RL deconvolution
        L = ||FP(X_all(ρ,nz),u)-W_u(ρ))||^2
        for l = 1:size(X,3) do
            I^(i)(ρ,lz) = I^(i)(ρ,(l-1)z)**G + I^(i)(ρ,(l+1)z)**G
            ∂L/∂F(l) = [2(FP(X_all(ρ,nz),u)-W_u(ρ))**H_u^T(ρ,nz)]
                          .* I^(i)(ρ,nz)
            // ** convolution operator
            // .* point multiplication operator
            F(l) = F(l) - α·∂L/∂F(l)
        end for
        // Update F using gradient descent algorithm
        for l = 1:size(X(,3) do
            I^(s)(ρ,lz) = I^(i)(ρ,lz) .* F(l)
        end for
        X(ρ,nz)  =  X_all(ρ,nz) - I^(s)(ρ,nz)
            // Update X(ρ,nz) based on X_all(ρ,nz)
    end for
end for
X(ρ,nz) = UP_xy(UP_z(X(ρ,nz),r_z),r_xy)
Return X(ρ,nz)
```

**Algorithm** RL_deconv

**Input:** point spread function of a specific spatial-angular component $H_u(\rho,nz)$, specific spatial-angular component $W_u(\rho)$, initial volume $X(\rho,nz)$, update rate $\lambda$

**Output:** updated 3D volume *Xguess*

```
HX(ρ) = FP(X(ρ,nz),u)
HXBack(ρ,nz) = BP(HX(ρ),u)
IBack(ρ,nz) = BP(W_u(ρ),u)
ErrorBack(ρ,nz) = IBack ./ HXBack
                    //  ./ point division operator
X(ρ,nz) = λ*X(ρ,nz)+(1-λ)*ErrorBack(ρ,nz)
Return X(ρ,nz)
```

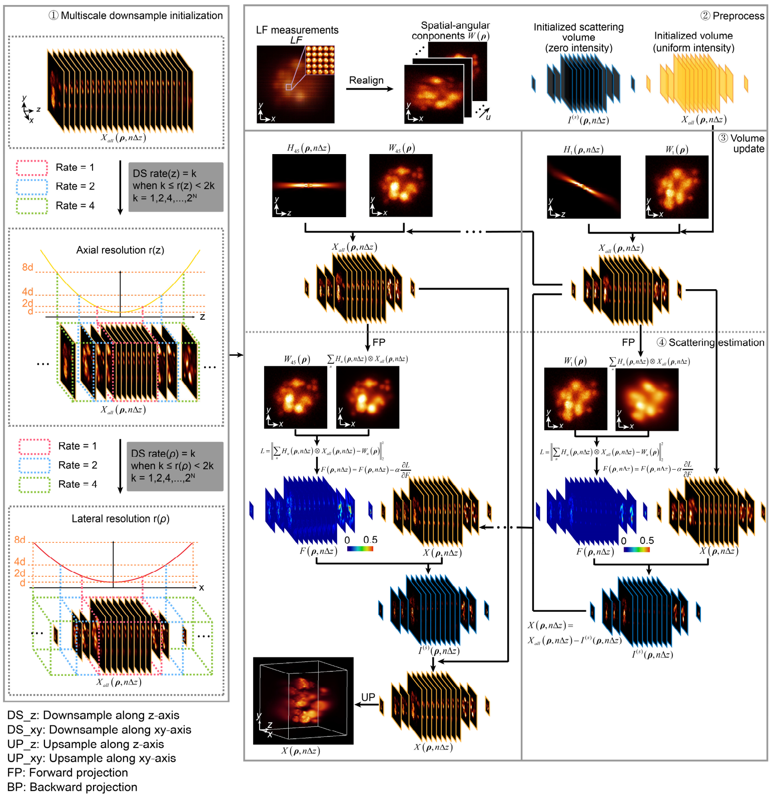


**Supplementary Figure 2 | The pseudo codes (left) and corresponding illustrations (right) of 3D deconvolution with a multiscale scattering model.** In traditional LF models, the volume initialized for reconstruction is densely sampled with a uniform sampling rate, which is very redundant due to the non-uniform low resolution of LFM at different axial planes. Consequently, only the high-resolution 3D range close to the native objective plane are modeled in traditional LFM to save the computational cost. On the contrary, our multiscale model considers a much larger volume to update the 3D non-uniform background fluorescence together with the high-resolution 3D range during reconstruction. Both the volume and point spread functions (PSF) are down-sampled with different sampling rates according to the characterized resolutions at different axial planes, resulting in orders-of-magnitude reduction in computational cost without resolution degradation. Then an ADMM framework is used to update the native 3D volume and scattered potentials iteratively. Only a specific angular component is used within each iteration to accelerate the convergence.

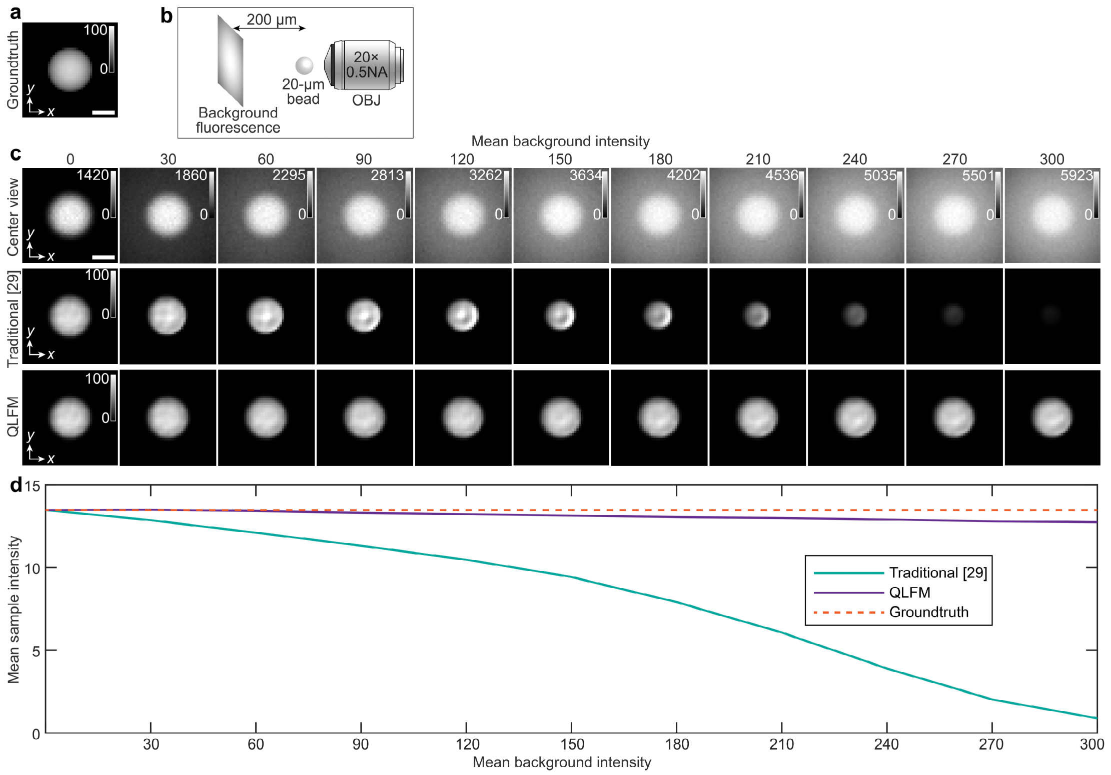


**Supplementary Figure 3 | Numerical analysis of the quantitative property of QLFM and traditional LFM with increasing background levels. a,** A simulated 20-μm-diameter fluorescence bead is placed at the focal plane as groundtruth with 100 photons in each voxel covering 0.33×0.33×0.5 μm$^3$. **b**, The simulated imaging schematic with a Gaussian-distributed background located at 200 μm away from the native objective plane. Both the background and the bead is imaged by the 20×/0.5NA objective. **c**, The simulated center views with shot noise and 42-μm mean intensity projections reconstructed by traditional LFM and QLFM, respectively, with different mean background intensities marked on the top. With the increase of the background, the structural information and the intensity will gradually decrease in traditional LFM, illustrating the loss of the quantitative property in deep tissue. On the contrary, QLFM show significantly-improved robustness against background fluorescence with the quantitative measurements. **d**, The curves of the mean intensities of the 27×27×42 μm$^3$ sub-volume reconstructed by different methods versus different mean background intensities, demonstrating the quantitative property of QLFM. The dashed line illustrates the groundtruth for comparison. Scale bar, 10 μm.

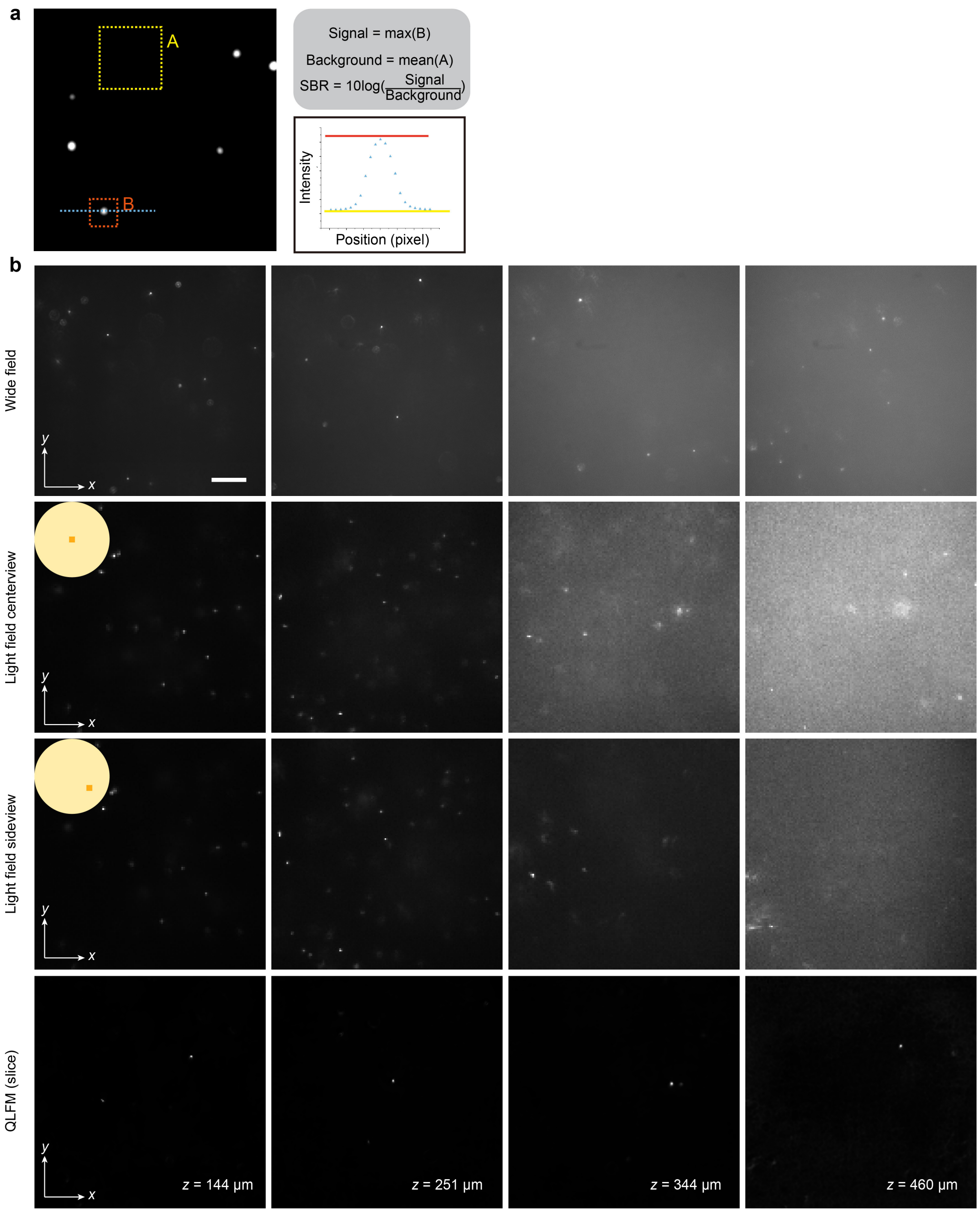


**Supplementary Figure 4 | Characterization of the signal-to-background ratio (SBR) in the tissue-mimicking phantom. a**, The SBR calculation. Average SBR is calculated by multiple fluorescence beads at the same penetration depth. **b**, We selected several typical areas at different penetration depths for comparisons among WFM, two angular views of LFM, and reconstructed slices by QLFM. QLFM show significantly-improved SBR over WFM, illustrating a larger penetration depth in deep tissue with computational optical sectioning. Scale bar, 100 μm.

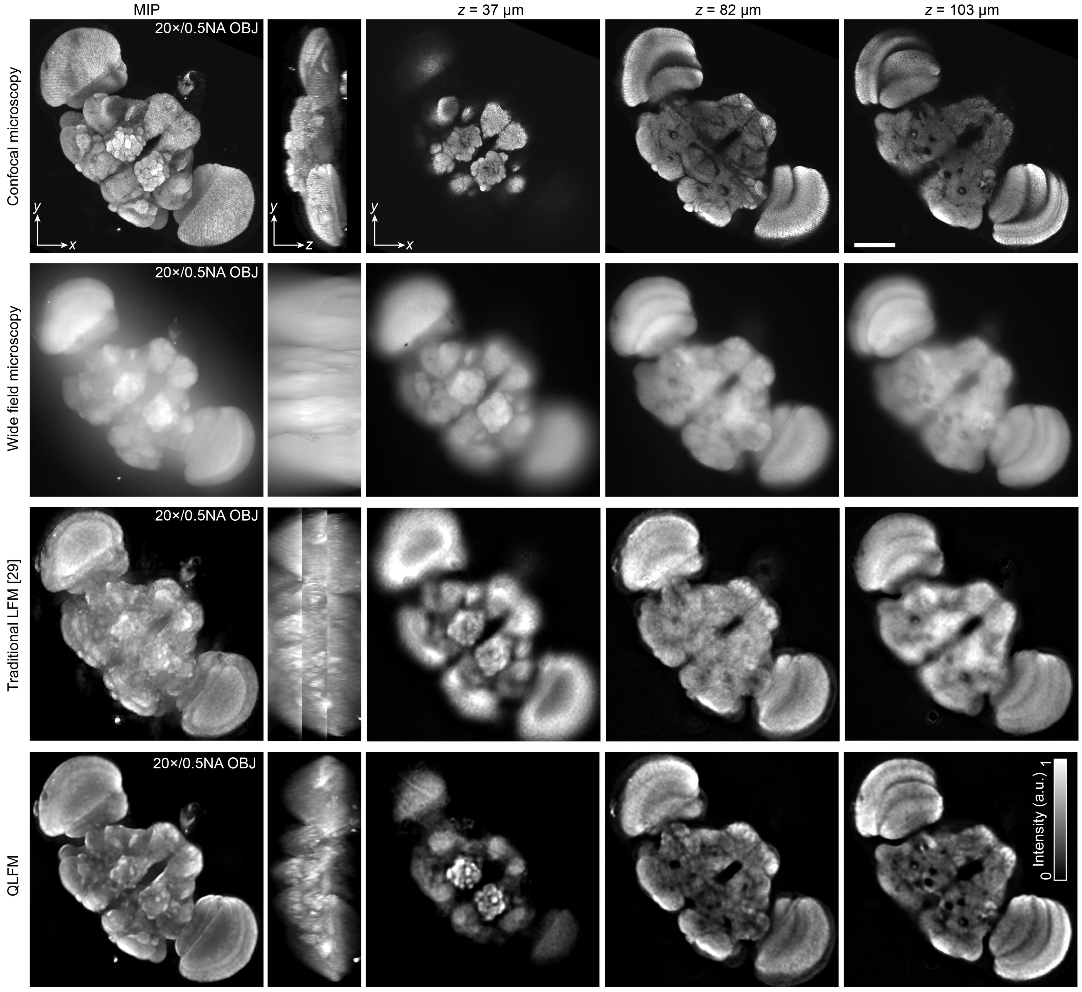


**Supplementary Figure 5 | Experimental comparisons among confocal microscopy, WFM, traditional LFM, and QLFM on the same fixed *Drosophila* brain.** The orthogonal MIPs and *xy*-slices obtained by different methods are shown on the different rows. During imaging, we used the same sample with a 20×/0.5NA objective. For confocal microscopy, we scanned the sample axially at a step of 1 μm for 139 slices. For WFM, we scanned 800 slices at 0.5-μm steps. For LFM, we scanned the entire volume with a step size of 50 μm by 3 times at 24 Hz, and reconstructed the volume by traditional LFM and QLFM, respectively. QLFM can not only get rid of the stitching artifacts with better contrast than traditional LFM, but also has a comparable optical-sectioning capability as confocal microscopy at a much faster 3D imaging speed. Scale bars, 100 μm.

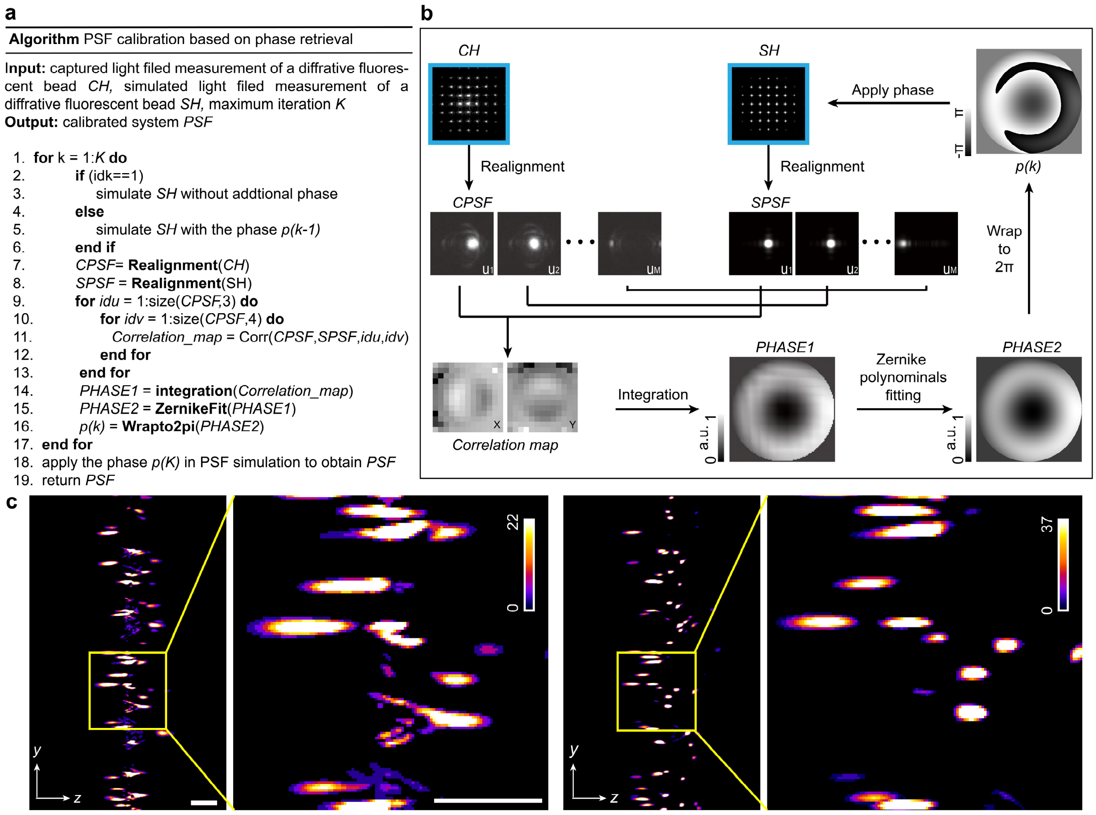


**Supplementary Figure 6 | The pseudo codes and corresponding illustrations for phase-retrieval-based PSF calibration.** **a**, Pseudo codes for the algorithm. **b**, Illustrations of the algorithm. As it's hard to capture the experimental PSF of LFM with a microlens array due to the high-dimensional property, simulated PSF based on wave optics is usually used during reconstruction. However, the system aberration of the experimental setup usually introduces severe system errors such as resolution degradation and artifacts. Here, we find that the accurate estimation of the PSF can eliminate the reconstruction artifacts close to the native objective plane in LFM and improve the high-resolution depth range. We used a single image of the sub-diffraction-limited fluorescence beads only, as marked by the blue box. Then, we generated the simulated LF PSF at a similar axial position. Both the captured and simulated images of a single point were realigned into phase-space domain in the form of multiple angular components, corresponding to different sub-apertures. Later, we calculated the disparities between the captured and simulated images along different angular components, which can be integrated as a phase map for calibration. After fitting with

Zernike polynomials, we apply the phase back to generate the PSF again for next iteration. The calibrated phase to estimate the system aberration will gradually converge after 5 iterations, which is then used for the calibrated PSF during reconstruction. **c**, Orthogonal MIPs of 0.5-μm fluorescence beads distributed in agarose reconstructed by QLFM with an ideal PSF without calibration (left) and a calibrated PSF (right), respectively. Scale bar, 20 μm.

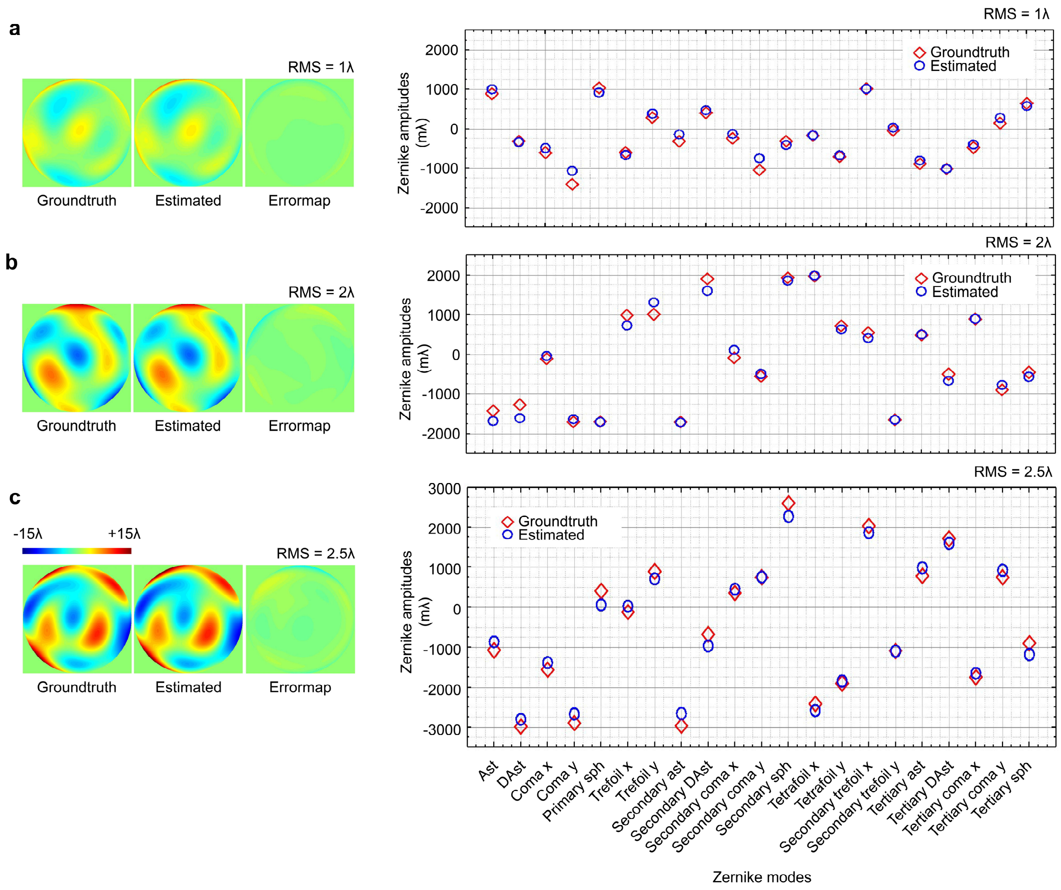


**Supplementary Figure 7 | Quantitative analysis of the phase-retrieval-based PSF calibration algorithm. a-c**, To evaluate the accuracy of the phase estimation, we conducted a numerical simulation with different levels of system aberrations applied to the system for a 20×/0.5NA objective. The groundtruth phase (left), the estimated phase by our algorithm (middle), and corresponding residual error map (right) are shown for comparisons. In addition, we visualized the amplitudes of 21 Zernike modes decomposed from the estimated pupils (blue circles) and the groundtruth (red diamonds), suggesting the effectiveness of the algorithm with a single image to calibrate the system.

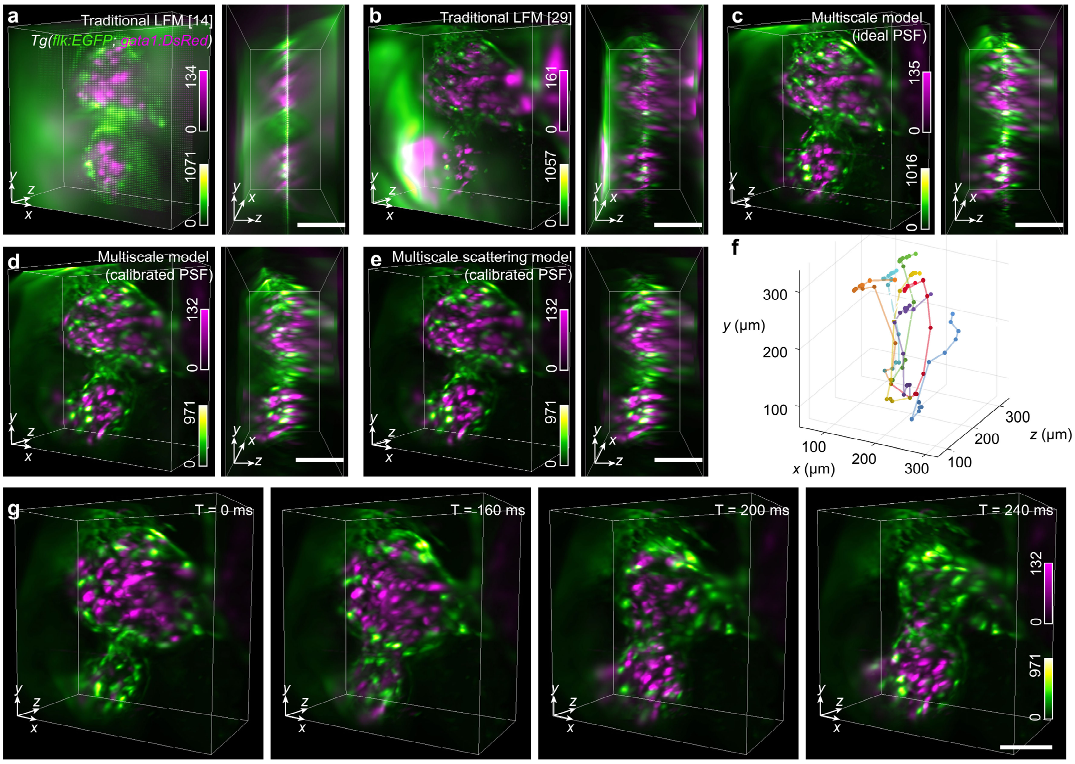


**Supplementary Figure 8 | Experimental comparisons on the beating heart in zebrafish larvae to show successive improvements. a-e**, The 3D rendered volumes reconstructed by traditional models, multiscale model with ideal PSFs, multiscale model with calibrated PSFs, and multiscale scattering model with calibrated PSFs, indicating the necessity of all the factors for quantitative volumetric reconstruction. The videos were captured with a 40×/1.0NA water-immersion objective at 25 Hz for two channels. All the methods are set to the same memory cost for fair comparisons. **f**, 3D tracking trajectories of representative blood cells flowing through the heart. **g**, 3D rendered volumes at different time stamps reconstructed by QLFM, showing engine-like dynamics of the heart-beating process. Scale bars, 50 μm

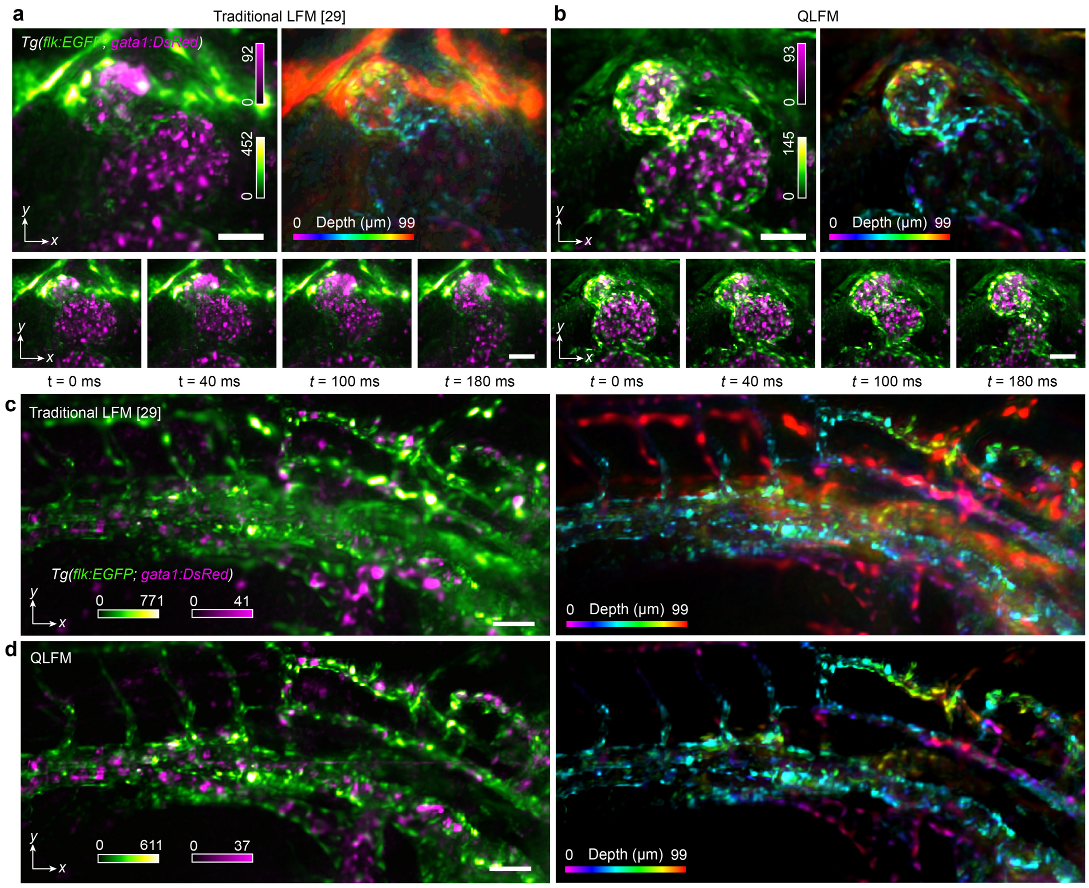


**Supplementary Figure 9 | Experimental comparisons on the large-scale blood flow in zebrafish larvae between traditional LFM and QLFM. a-b**, The two-color MIPs and depth-coded MIPs of a beating heart in zebrafish larvae reconstructed by traditional LFM and QLFM. The two-color MIPs at different time stamps are marked at the bottom row, illustrating the large-scale red blood cells (magenta) flowing in the vessels (green). For the depth-coded MIP, different colors correspond to different depths. **c-d**, The orthogonal MIPs and depth-coded MIPs for comparison in another region, demonstrating the application of large-scale 3D tracking. Scale bars, 50 μm.

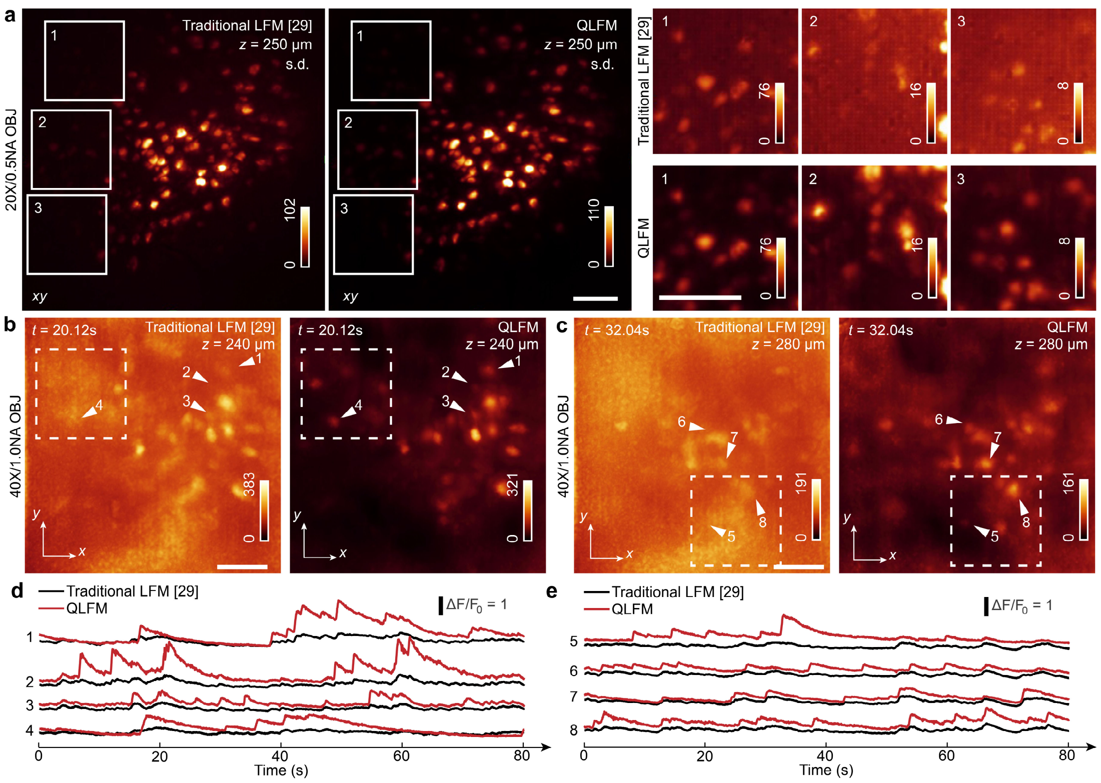


**Supplementary Figure 10 | *In vivo* 3D calcium imaging in mouse cortex. a**, The MIPs of the standard deviations (s.d.) for the 2000 volumes obtained by traditional LFM and QLFM under a 20×/0.5NA objective (Supplementary Video 4). The MIPs are displayed at different contrast due to the high dynamic range of QLFM, indicating that QLFM can detect more neurons due to improved SBR. **b-c**, The comparisons of the MIPs at *t*=20.12s between traditional LFM and QLFM under a 40×/1.0NA water-immersion objective imaged at different depths, showing QLFM has larger penetration depth with computational optical sectioning. **d-e,** Temporal traces of marked neurons in b and c, illustrating the better contrast. Scale bars, 100 μm (**a**) and 50 μm (**b-c**).

### Supplementary Notes

#### Supplementary Note 1. Incoherent scattering model for 3D deconvolution.

Considering a monochromatic electromagnetic field (the time-dependent factor $\exp(-\pi\omega t)$ is not shown), the scattering medium is limited to a finite large area $V$ modeled in a multiscale way, and satisfies linearity, isotropy and non-magnetic property. The space-dependent part of the complex electric field $\mathbf{E}$ can be expressed by:

$$\nabla^2\mathbf{E}(\mathbf{r},\omega)+k^2\varepsilon(\mathbf{r},\omega)\mathbf{E}(\mathbf{r},\omega)+\mathrm{grad}[\mathbf{E}(\mathbf{r},\omega)\cdot\mathrm{grad}\ln\varepsilon(\mathbf{r},\omega)]=0\,, \tag{1}$$

where $\mathbf{r}=(x,y,z)$ is the 3D spatial positions, $\omega$ is the frequency coordinate, $k=\omega/c$ is the wave number, and $\nabla^2$ is the Laplace operator. Then, we assume the dielectric coefficient $\varepsilon(\mathbf{r},\omega)$ is approximately constant in the range of center wavelength $\lambda=2\pi/k=2\pi(c/\omega)$ with the emission filter. So, Eq. (1) can be simplified as

$$\nabla^2\mathbf{E}(\mathbf{r},\omega)+k^2n^2(\mathbf{r},\omega)\mathbf{E}(\mathbf{r},\omega)=0\,, \tag{2}$$

where $n(\mathbf{r},\omega)$ is the medium refractive index, satisfying Maxwell's equation $\varepsilon(\mathbf{r},\omega)=n^2(\mathbf{r},\omega)$. The scalar form of Eq. (2) is written as:

$$\nabla^2\mathbf{U}(\mathbf{r},\omega)+k^2n^2(\mathbf{r},\omega)\mathbf{U}(\mathbf{r},\omega)=0\,. \tag{3}$$

The Eq. (3) is tranformed into

$$\begin{gathered}\nabla^2U(\mathbf{r},\omega)+k^2U(\mathbf{r},\omega)=-4\pi F_0(\mathbf{r},\omega)U(\mathbf{r},\omega)\\ F_0(\mathbf{r},\omega)=\frac{1}{4\pi}k^2\left[n^2(\mathbf{r},\omega)-1\right]\end{gathered}\,, \tag{4}$$

where $F_0(\mathbf{r},\omega)$ is defined as the scattering potential energy. The field $U(\mathbf{r},\omega)$ represents the sum of incident field $U^{(i)}(\mathbf{r},\omega)$ and scattering field $U^{(s)}(\mathbf{r},\omega)$:

$$U(\mathbf{r},\omega)=U^{(i)}(\mathbf{r},\omega)+U^{(s)}(\mathbf{r},\omega)\,. \tag{5}$$

The incident field usually can be regarded as a plane wave, which satisfies Helmholtz equation:

$$\left(\nabla^2 + k^2\right)U^{(i)}(\mathbf{r},\omega) = 0 \cdot \tag{6}$$

The intensity field can be obtained as:

$$I(\mathbf{r},\omega) = U(\mathbf{r},\omega)\cdot U^*(\mathbf{r},\omega). \tag{7}$$

By substituting Eq.(5) into Eq.(7) and ignoring the crossed terms, the intensity field is written as:

$$I(\mathbf{r},\omega) = I^{(i)}(\mathbf{r},\omega) + I^{(s)}(\mathbf{r},\omega)\,, \tag{8}$$

where $I^{(i)}(\mathbf{r},\omega)$ is the incident intensity and $I^{(s)}(\mathbf{r},\omega)$ is the scattering intensity. The second order derivative of the intensity field $I(\mathbf{r},\omega)$ is then developed into:

$$\begin{aligned}\nabla^2 I(\mathbf{r},\omega) &= \nabla^2(U(\mathbf{r},\omega)\cdot U^*(\mathbf{r},\omega)) \\ &= U(\mathbf{r},\omega)\nabla^2 U^*(\mathbf{r},\omega) + U^*(\mathbf{r},\omega)\nabla^2 U(\mathbf{r},\omega)\end{aligned}. \tag{9}$$

Using Eq. (6), Eq. (9) can be tranformed into:

$$\nabla^2 I(\mathbf{r},\omega) + 2k^2 I(\mathbf{r},\omega) = -4\pi F(\mathbf{r},\omega)I(\mathbf{r},\omega)\,, \tag{10}$$

where

$$F(\mathbf{r},\omega) = 2F_0(\mathbf{r},\omega) = \frac{1}{2\pi}k^2\left[n^2(\mathbf{r},\omega) - 1\right]. \tag{11}$$

Using Eq. (6) and Eq. (8), Eq. (10) can be tranformed into:

$$(\nabla^2 + 2k^2)I^{(s)}(\mathbf{r},\omega) = -4\pi F(\mathbf{r},\omega)I(\mathbf{r},\omega)\,. \tag{12}$$

Then, we convert the differential equation to an integral equation. We let $G(\mathbf{r}-\mathbf{r}')$ be the Green function of Helmholtz operator, which satisfies

$$\left(\nabla^2 + k^2\right)G\left(\mathbf{r}-\mathbf{r}',\omega\right) = -4\pi\delta^{(3)}\left(\mathbf{r}-\mathbf{r}'\right), \tag{13}$$

where $\delta^{(3)}(\cdot)$ is the 3D Dirac impulse function. We take the conjugate of Eq. (13) to obtain

$$\left(\nabla^2 + k^2\right)G^*\left(\mathbf{r}-\mathbf{r}',\omega\right) = -4\pi\delta^{(3)}\left(\mathbf{r}-\mathbf{r}'\right)\cdot \tag{14}$$

Eq. (13) and Eq. (14) can be combined as:

$$\left(\nabla^2+2k^2\right)\left\|G\left(\mathbf{r}-\mathbf{r}',\omega\right)\right\|_2^2=-4\pi\delta^{(3)}\left(\mathbf{r}-\mathbf{r}'\right)\left[G\left(\mathbf{r}-\mathbf{r}',\omega\right)+G^*\left(\mathbf{r}-\mathbf{r}',\omega\right)\right]. \quad (15)$$

Then with Eq. (11), we can obtain:

$$\begin{aligned}&I^{(s)}(\mathbf{r},\omega)\nabla^2\left\|G\left(\mathbf{r}-\mathbf{r}',\omega\right)\right\|_2^2-\left\|G\left(\mathbf{r}-\mathbf{r}',\omega\right)\right\|_2^2\nabla^2 I^{(s)}(\mathbf{r},\omega)\\&=4\pi F(\mathbf{r},\omega)I(\mathbf{r},\omega)\left\|G\left(\mathbf{r}-\mathbf{r}',\omega\right)\right\|_2^2\\&\quad-4\pi I^{(s)}(\mathbf{r},\omega)\delta^{(3)}\left(\mathbf{r}-\mathbf{r}'\right)\left[G\left(\mathbf{r}-\mathbf{r}',\omega\right)+G^*\left(\mathbf{r}-\mathbf{r}',\omega\right)\right]\end{aligned}. \quad (16)$$

By converting volume integration to surface integration, we have

$$\begin{aligned}&I^{(s)}(\mathbf{r},\omega)\\&=\frac{1}{2}\int_V F\left(\mathbf{r}',\omega\right)I\left(\mathbf{r}',\omega\right)\left\|G\left(\mathbf{r}-\mathbf{r}',\omega\right)\right\|_2^2\mathrm{d}^3r'\\&=-\frac{1}{8\pi}\int_{S_R}\left[I^{(s)}\left(\mathbf{r}',\omega\right)\frac{\partial\left\|G\left(\mathbf{r}-\mathbf{r}',\omega\right)\right\|_2^2}{\partial n'}-\left\|G\left(\mathbf{r}-\mathbf{r}',\omega\right)\right\|_2^2\frac{\partial I^{(s)}\left(\mathbf{r}',\omega\right)}{\partial n'}\right]\mathrm{d}S_R\end{aligned}, \quad (17)$$

where $V$ is the range of volume integration, and the spherical surface $S_R$ surrounds the scattering area[33]. Here, we select the Green function as below (from now on we leave out the frequency $\omega$):

$$G\left(\mathbf{r}-\mathbf{r}'\right)=\frac{\mathrm{e}^{\mathrm{i}k\left|\mathbf{r}-\mathbf{r}'\right|}}{\left|\mathbf{r}-\mathbf{r}'\right|}, \quad (18)$$

and

$$\left\|G\left(\mathbf{r}-\mathbf{r}'\right)\right\|_2^2=\frac{1}{\left\|\mathbf{r}-\mathbf{r}'\right\|_2^2}. \quad (19)$$

In the far-field range of the scattering volume, the scattered field can often be assumed to have the characteristics of a spherical wave. Taking the limit $R\to\infty$, the surface integral on the right side of Eq. (17) has no contribution to the total field[43,44]. Therefore, the scattered field is simplified as:

$$I^{(s)}(\mathbf{r})=\frac{1}{2}\int_V F\left(\mathbf{r}'\right)I\left(\mathbf{r}'\right)\frac{1}{\left\|\mathbf{r}-\mathbf{r}'\right\|_2^2}\mathrm{d}^3r'. \quad (20)$$

For simplicity, the total field in Eq. (20) is approximated as the incident intensity based on the assumption of weak scattering, leading to the expression as below:

$$I^{(s)}(\mathbf{r})=\frac{1}{2}\int_V F\left(\mathbf{r}'\right)I^{(i)}\left(\mathbf{r}'\right)\frac{1}{\left\|\mathbf{r}-\mathbf{r}'\right\|_2^2}\mathrm{d}^3r'. \quad (21)$$

where $\mathbf{r}$ can be written as $\mathbf{r}=(\boldsymbol{\rho},z)$, $\boldsymbol{\rho}=(x,y)$ is the lateral coordinate, and $z$ is the axial coordinate.

For the discrete model in reconstruction, the 3D object is divided into multiple slices. Each slice has a thin but finite thickness $\Delta z$. The relation between the incident intensity and the scattering intensity in $n^{\text{th}}$ slice can be written as:

$$\begin{aligned} I^{(s)}(\boldsymbol{\rho},n\Delta z) &= \frac{1}{2}\Delta z\int_S F\left(\boldsymbol{\rho}^{'},n\Delta z\right)I^{(i)}\left(\boldsymbol{\rho}^{'},n\Delta z\right)\frac{1}{\left\|\boldsymbol{\rho}-\boldsymbol{\rho}^{'}\right\|_2^2}\mathrm{d}\boldsymbol{\rho}^{'} \\ &= C\left[F\left(\boldsymbol{\rho},n\Delta z\right)I^{(i)}\left(\boldsymbol{\rho},n\Delta z\right)\right]\otimes\frac{1}{\left\|\boldsymbol{\rho}\right\|_2^2} \\ &= C\left[F\left(\boldsymbol{\rho},n\Delta z\right)I^{(i)}\left(\boldsymbol{\rho},n\Delta z\right)\right]\otimes\delta\left(\boldsymbol{\rho}\right) \\ &= CF\left(\boldsymbol{\rho},n\Delta z\right)I^{(i)}\left(\boldsymbol{\rho},n\Delta z\right) \end{aligned}, \qquad (22)$$

where $\delta(\cdot)$ is the 2D Dirac impulse function, $C$ is a constant, omitted for simplicity, $S$ is the range of one slice, and $\otimes$ represents the convolution operator. Considering forward and backward scattering processes, the incident intensity in $n^{\text{th}}$ slice can be divided as:

$$I^{(i)}\left(\boldsymbol{\rho},n\Delta z\right)=I_F^{(i)}\left(\boldsymbol{\rho},n\Delta z\right)+I_B^{(i)}\left(\boldsymbol{\rho},n\Delta z\right), \qquad (23)$$

where $I_F^{(i)}\left(\boldsymbol{\rho},n\Delta z\right)$ represents the forward incident intensity in $\text{n}^{\text{th}}$ slice and $I_B^{(i)}\left(\boldsymbol{\rho},n\Delta z\right)$ represents the backward incident intensity. These formulas can be written as:

$$\begin{aligned} I_F^{(i)}\left(\boldsymbol{\rho},n\Delta z\right) &= \left[I_F^{(i)}\left(\boldsymbol{\rho},(n-1)\Delta z\right)+X\left(\boldsymbol{\rho},(n-1)\Delta z\right)\right]\otimes\frac{1}{\left\|\mathbf{r}\right\|_2^2} \\ I_B^{(i)}\left(\boldsymbol{\rho},n\Delta z\right) &= \left[I_B^{(i)}\left(\boldsymbol{\rho},(n+1)\Delta z\right)+X\left(\boldsymbol{\rho},(n+1)\Delta z\right)\right]\otimes\frac{1}{\left\|\mathbf{r}\right\|_2^2}, \\ I_B^{(i)}\left(\boldsymbol{\rho},N\Delta z\right) &= 0 \\ I_F^{(i)}\left(\boldsymbol{\rho},0\right) &= 0 \end{aligned} \qquad (24)$$

where $X\left(\boldsymbol{\rho},n\Delta z\right)$ represents the emitted fluorescence intensity in $n^{\text{th}}$ slice and $N$ is the total number of the slices. The Eq. (22) can be developed into

$$\begin{aligned} I^{(s)}(\boldsymbol{\rho}, n\Delta z) &= F(\boldsymbol{\rho}, n\Delta z) I^{(i)}(\boldsymbol{\rho}, n\Delta z) \\ &= F(\boldsymbol{\rho}, n\Delta z) \cdot \left[ I_F^{(i)}(\boldsymbol{\rho}, n\Delta z) + I_B^{(i)}(\boldsymbol{\rho}, n\Delta z) \right] \end{aligned}. \tag{25}$$

To estimate the scattering intensity and the fluorescence intensity, we consider $I^{(s)}(\mathbf{r}, \omega)$ in the 3D deconvolution algorithm with ADMM framework[34], as shown in Supplementary Fig. 2. The whole 3D volume and PSF are sampled according to the designed multiscale sampling rate. Within each iteration, the phase-space deconvolution is used to update the reconstructed volume $X_{all}(\boldsymbol{\rho}, n\Delta z)$, which consists of the native fluorescence intensity $X(\boldsymbol{\rho}, n\Delta z)$ and the scattering intensity $I^{(s)}(\boldsymbol{\rho}, n\Delta z)$:

$$X_{all}(\boldsymbol{\rho}, n\Delta z) = X(\boldsymbol{\rho}, n\Delta z) + I^{(s)}(\boldsymbol{\rho}, n\Delta z). \tag{26}$$

The $I^{(s)}(\boldsymbol{\rho}, n\Delta z)$ and $X(\boldsymbol{\rho}, n\Delta z)$ can then be obtained directly from the fixed scattering potential $F(\boldsymbol{\rho}, n\Delta z)$ based on Eq. (25). $F(\boldsymbol{\rho}, n\Delta z)$ is initialized to zero for the first iteration.

Then, we fix the sample intensity $X(\boldsymbol{\rho}, n\Delta z)$ to update the scattering potential $F(\boldsymbol{\rho}, n\Delta z)$ with the gradient descent method to solve the following optimization problem:

$$\begin{aligned} &\min_F L \\ L &= \left\| \sum_n H_u(\boldsymbol{\rho}, n\Delta z) \otimes X_{all}(\boldsymbol{\rho}, n\Delta z) - W_u(\boldsymbol{\rho}) \right\|_2^2 \\ &= \left\| \sum_n H_u(\boldsymbol{\rho}, n\Delta z) \otimes \left\{ X(\boldsymbol{\rho}, n\Delta z) + F(\boldsymbol{\rho}, n\Delta z) I^{(i)}(\boldsymbol{\rho}, n\Delta z) \right\} - W_u(\boldsymbol{\rho}) \right\|_2^2 \\ &\text{s.t. } 0 \le F(\boldsymbol{\rho}, n\Delta z) \le 1 \end{aligned}, \tag{27}$$

where $H_u(\boldsymbol{\rho}, n\Delta z)$ is the PSF for a specific angle $u$, and $W_u(\boldsymbol{\rho})$ is the captured angular component along angle $u$ after pixel realignemnt. Then the update formula for the scattering potential $F(\boldsymbol{\rho}, n\Delta z)$ can be written as :

$$F(\boldsymbol{\rho}, n\Delta z) = F(\boldsymbol{\rho}, n\Delta z) - \alpha \frac{\partial L}{\partial F}, \tag{28}$$

where $\frac{\partial L}{\partial F} = 2\left(\sum_n H_u(\boldsymbol{\rho}, n\Delta z) \otimes X_{all}(\boldsymbol{\rho}, n\Delta z) - W_\omega(\boldsymbol{\rho})\right) \otimes H_u{}^T(-\boldsymbol{\rho}, n\Delta z) \cdot I^{(i)}(\boldsymbol{\rho}, n\Delta z)$, $H_u{}^T(-\boldsymbol{\rho}, n\Delta z)$ is the transpose PSF for a specific angle and $\alpha$ is the learning rate. Then, the updated scattering potential $F(\boldsymbol{\rho}, n\Delta z)$ is fed again to the next iteration for another angle. We repeat the above ADMM steps along every angular components iteratively for convergence. Each angular component is used for only one time during iterations for all the experiments without axial scanning, while each angular component is used for two times for all the experiments with axial scanning.

**References**


[33] Born, M., Wolf, E. & Hecht, E. Principles of Optics: Electromagnetic Theory of Propagation, Interference and Diffraction of Light. *Phys. Today* (2000).

[34] Boyd, Stephen, Neal Parikh, Eric Chu, Borja Peleato, and Jonathan Eckstein. "Distributed Optimization and Statistical Learning via the Alternating Direction Method of Multipliers." *Foundations and Trends in Machine Learning*. (2010).

[43] Colton, David, and Rainer Kress. Inverse Acoustic and Electromagnetic Scattering Theory: Fourth Edition. *Applied Mathematical Sciences*. (2019).

[44] Sneddon, I. N., B. B. Baker, and E. T. Copson. "The Mathematical Theory of Huygens' Principle." *The Mathematical Gazette*. (1951).

**Supplementary Table 1. Imaging conditions for all fluorescence experiments.**

| | Sample, (imaging T, °C) | Fluorescent label | Exposure time (# time pts) | λ: Power ($mW/mm^2$) | Volume rate (Hz) | Objective | Axially-scanned planes and spacing |
|---|---|---|---|---|---|---|---|
| 1c, 1d, S2, SV1 | Tumor spheroid 27 °C | GFP | 100 ms 1 pts | 488: 1.21 | - | 63×/1.4NA Oil | 10 planes 10 μm |
| 2b, S4 | Tissue-mimicking phantom 27 °C | Yellow-green fluorescent (505/515) | 20 ms 800 pts | 488: 11.7 | - | 40×/1.0NA Water | 1 plane |
| 2c | Fluorescence beads 27 °C | Yellow-green fluorescent (505/515) | 95 ms 1 pts | 488: 11.7 | - | 40×/1.0NA Water | 1 plane |
| 2d, S6c | Fluorescence beads 27 °C | Yellow-green fluorescent (505/515) | 100 ms 100 pts | 488: 11.7 | - | 40×/1.0NA Water | 5 planes 30 μm |
| 2g | Zebrafish larval 27°C | *Tg(flk:EGFP; gata1:DsRed)* | 10 ms 1 pts | 488: 11.2 561: 8.1 | - | 20×/0.5NA Air | 1 plane |
| 3a-b | *Drosophila* embryos 25℃ | EGFP (His2Av) | 95 ms 1 pts | 488: 11.7 | - | 40×/1.0NA Water | 8 planes 15 μm |
| 3c-d | *Drosophila* embryos 25℃ | EGFP (His2Av) | 300 ms 1600 pts | 488: 7.1 | 1 | 20×/0.5NA Air | 3 planes 50 μm |
| 4a-c, S8, SV2 | Zebrafish larval 27°C | *Tg(flk:EGFP; gata1:DsRed)* | 10 ms 100 pts | 488: 61.9 561:80.5 | 25 | 40×/1.0NA Water | 1 plane |
| 4d-g, SV3 | Zebrafish larval 27°C | *Tg(huc:GCamp6)* | 30 ms 4000 pts | 488: 12.5 | 24 | 20×/0.5NA Air | 1 plane |

| | | | | | | | |
|---|---|---|---|---|---|---|---|
| 5a-c, S10a SV4 | Awake behaving mouse 27℃ | GCaMP6s | 30 ms 2000 pts | 488:3.6 | 25 | 20×/0.5NA Air | 1 plane |
| 5d-e, S10b-e, SV5 | Awake behaving mouse 27℃ | GCaMP6s | 30 ms 2000 pts | 488:3.4 | 25 | 40×/1.0NA Water | 1 plane |
| 5f, SV6 | Awake behaving mouse 27℃ | GCaMP6f | 30 ms 500 pts | 488: 1.4 | 6 | 20×/0.5NA Air | 3 planes 50 μm |
| S5 | *Drosophila* brain 25℃ | Antibody nc82 | 100 ms 1 pts | 488: 11.0 | - | 20×/0.5NA Air | 5 planes 50 μm |
| S9 | Zebrafish larval 27°C | *Tg(flk:EGFP; gata1:DsRed)* | 10 ms 50 pts | 488: 5.0 561: 9.0 | 25 | 20×/0.5NA Air | 1 plane |
